# Different temporal requirements for the LRR transmembrane receptors Tartan and Toll-2 in the formation of contractile interfaces at compartmental boundaries

**DOI:** 10.1101/2021.12.17.473176

**Authors:** Thomas E. Sharrock, Guy B. Blanchard, Jenny Evans, Bénédicte Sanson

## Abstract

Compartmental boundaries physically separate groups of epithelial cells, a property fundamental for the organization of the body plan in both insects and vertebrates. In many examples, this physical separation has been shown to be the consequence of a regulated increase in contractility of the actomyosin cortex at boundary cell-cell interfaces, a property important in developmental morphogenesis beyond compartmental boundary formation. In this study, we took an unbiased screening approach to identify cell surface receptors required for actomyosin enrichment and polarisation at parasegmental boundaries (PSBs) in early *Drosophila* embryos, leading us to uncover different temporal requirements for two LRR receptors, Tartan and Toll-2. First, we find that Tartan is required during germband extension for actomyosin enrichment at PSBs, confirming an earlier report. Next, by following in real time the dynamics of loss of boundary straightness in *tartan* mutant embryos compared to wildtype and *ftz* mutant embryos, we show that Tartan is not required beyond germband extension. At this stage, actomyosin enrichment at PSBs becomes regulated by Wingless signalling. We find that Wingless signalling regulates Toll-2 expression and we show that Toll-2 is required for planar polarization of actomyosin after the completion of germ-band extension. Thus the formation of contractile interfaces at PSBs depends on a dynamic set of LRR receptors cues. Our study also suggests that the number of receptor cues is small and that the receptors are interchangeable.

## Introduction

The mechanisms underlying the partitioning group of cells into immiscible compartments have fascinated scientists since the discovery of compartmental boundaries in *Drosophila* in the 1970s (Fagotto, 2020a). In many cases studied, this physical barrier is caused by a localised upregulation of actomyosin contractility at boundary cell-cell interfaces, found in both *Drosophila* and vertebrate models (Aliee et al., 2012; Calzolari et al., 2014; Canty et al., 2017; Landsberg et al., 2009; Monier et al., 2010). How this increase in cortex contractility is specified at boundary interfaces to create mechanical barriers remains only partially understood. Within homogeneous fields of epithelial cells, spatial regulation of transcription factors is key to initiate boundary formation (Dahmann et al., 2011; Monier et al., 2011). Downstream of these transcription factors, various cell surface receptors have been implicated in causing actomyosin enrichment at boundary interfaces. In vertebrates, for example at rhombomere boundaries in the hindbrain, the Ephrin/Eph receptors play a key role, but additional cell surface asymmetries have also been identified (Fagotto, 2020b; Pujades, 2020). In *Drosophila*, downstream receptors remained elusive for a long time, but recent work is starting to identify specific cell surface asymmetries required for the formation of mechanical boundaries (Sharrock and Sanson, 2020; Wang and Dahmann, 2020).

Beyond its role in compartmental cell sorting, increase in cortical contractility at epithelial cell-cell junctions underlies many cell and tissue behaviours (Amack and Manning, 2012; Bielmeier et al., 2016; Bosveld et al., 2016; Collinet and Lecuit, 2021). In particular, convergent extension, where cells intercalate to elongate a tissue, was shown in *Drosophila* to require planar polarized enrichment of the actomyosin cortex (Bertet et al., 2004; Collinet and Lecuit, 2021; Pare and Zallen, 2020; Zallen and Wieschaus, 2004). This planar polarization is downstream of antero-posterior patterning, which generates the subdivision of the body axis by overlapping stripes of transcription factors encoded by the pair-rule genes (Bertet et al., 2004; Irvine and Wieschaus, 1994; Zallen and Wieschaus, 2004). Actomyosin planar polarization has now been linked to convergent extension in several vertebrate examples, and is generally thought to be regulated by the Planar Cell Polarity (PCP) pathway (Collinet and Lecuit, 2021; Nishimura et al., 2012; Pare and Zallen, 2020; Rozbicki et al., 2015; Shindo and Wallingford, 2014). It is not known, however, whether gene expression asymmetries contribute to the activation of the PCP pathway to drive convergent extension in vertebrates.

In this paper, we have searched for cell surface receptors driving actomyosin enrichment at compartmental boundaries, using an unbiased screening approach. We focused on parasegmental boundaries that subdivide the germband in early *Drosophila* embryos. Parasegmental boundaries (PSBs) form during convergent extension of the germband during gastrulation, a process called germband extension (GBE). We had shown previously that actomyosin enrichments form at PSBs in the course of GBE, progressively emerging from the tissue-wide actomyosin planar polarization that is initiated at gastrulation (Tetley et al., 2016). Tissue-wide planar polarization of actomyosin requires the LRR receptors Toll-like 2, 6 and 8, which are expressed in overlapping stripes downstream of the pair-rule transcriptional network (Pare et al., 2014). However, removal of all three receptors is not sufficient to abolish actomyosin enrichment at PSBs, suggesting that additional receptor(s) are required (Pare et al., 2019; Pare et al., 2014). In (Tetley et al., 2016), modeling cell-cell interactions during GBE, we proposed that an additional receptor expressed in a periodic, double-segment pattern, might be sufficient to confer polarization of actomyosin at PSBs. We undertook a systematic screen based on this hypothesis, which we present here. A second question concerned the role of surface receptors at different developmental stages. Once axis extension is completed, actomyosin enrichments are maintained at PSBs during the extended germband stages (Monier et al., 2010). Whereas actomyosin enrichments during convergent extension (GBE, stages 7-8) require the pair-rule gene network, their maintenance at PSBs after completion of germband extension (stages 9-10-11) requires Wingless signalling (Monier et al., 2010; Scarpa et al., 2018; Urbano et al., 2018). No cell surface receptors had yet been identified downstream of Wingless signalling, so we also addressed this as part of our screening approach.

From our screen we find that Tartan, another LRR receptor, is required for actomyosin enrichment at PSBs throughout axis extension. This provides an independent confirmation of earlier findings by (Pare et al., 2019). By visualising transcription of parasegmental markers in combination with cell tracking in live embryos, we are able to follow how boundary straightness (a functional consequence of actomyosin enrichment at boundaries) evolves in the course of axis extension in wild-type and *tartan* mutant embryos and also in the pair-rule mutant *ftz*, in which *tartan* anteroposterior stripes are lost. This analysis showed that *tartan* is required for for specifiying contractile interfaces at PSB during GBE, but not beyond. Rather, we show that the expression of the LRR receptor Toll-2 is regulated by Wingless signalling and is required for actomyosin enrichment at PSBs during the extended germband stages. This work shows that a temporal succession of LRR receptor asymmetries specify planar polarized mechanical interfaces at PSBs.

## Results

### A screen to find cell surface receptors expressed asymmetrically at parasegmental boundaries

From our previous work on axis extension, we predicted that the expression of a single surface molecule within either even or odd-numbered parasegments would constitute the minimal requirement for generating the missing molecular asymmetries at the parasegmental boundary (Tetley et al., 2016). Based on this prediction, we performed an *in silico* screen to find genes meeting the following three criteria: i) they should be expressed in stripes along the anteroposterior (AP) axis ii) they should encode a protein that localizes to the cell surface and iii) they should be regulated by the pair-rule gene network. Mining publicly available data for the 13,600 genes in the *Drosophila* genome, we identified 822 genes expressed in AP stripes, 5620 genes encoding proteins with a signal peptide and/or a predicted transmembrane domain and 3679 genes likely to be pair-rule regulated (Methods). Following standardization of the gene nomenclature using the unique Flybase Identifiers, we found 94 genes in common with these three datasets (Figure 1A). Next, we applied manual quality controls to this initial list, whittling the number of candidates down to 31 genes (Methods). Excluded genes were those not showing an obvious stripy pattern by eye in *in situ* libraries or being likely to be expressed at very low levels in early embryos or having a known localization in the literature different from the cell surface (e.g. transcription factors) (Figure 1B and Methods).

**Figure 1:**
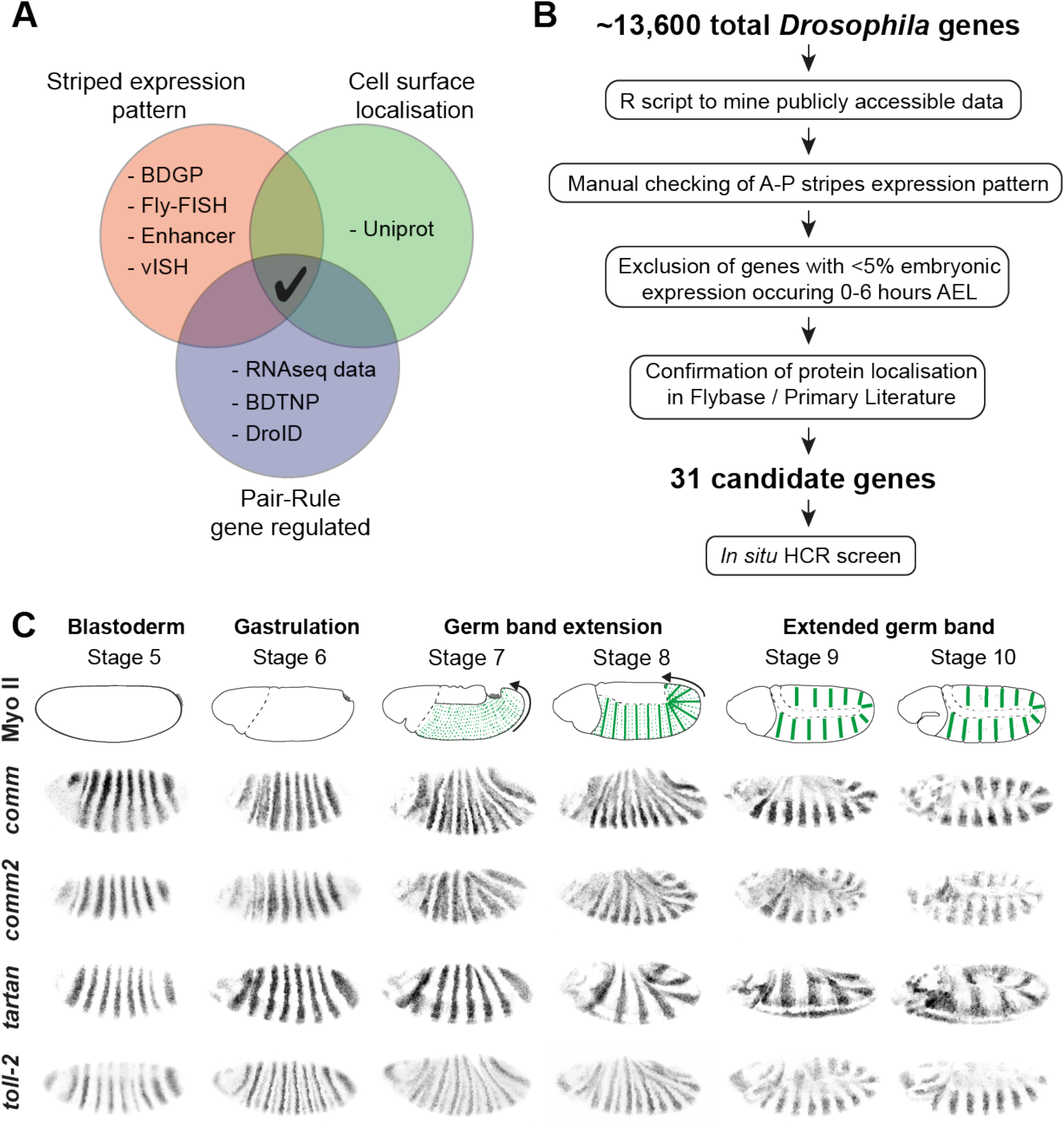
*In silico* screen to identify cell surface factors responsible for actomyosin enrichment at PSBs. A) Venn diagram illustrating the three criteria used to identify candidate cell surface factors in an *in silico* screen and including the main datasets that were mined. B) Flow chart showing the successive steps taken to whittle down candidates. C) *In situ* HCR images for key candidate genes found in the screen. *comm, comm2, tartan* and *toll-2* are all expressed in clear anteroposterior stripes in early embryos. We distinguish two developmental periods for actomyosin enrichment at parasegmental boundaries (PSBs), germband extension (GBE) and extended germband stages. At the onset of GBE, actomyosin becomes planar polarised at every AP cell-cell interfaces, including PSB interfaces (dashed green lines, stage 7). As GBE progresses and cells intercalate (stage 8), actomyosin enrichment become more prominent at PSBs (green lines). Weaker actomyosin enrichment is also detectable at two intraparasegmental boundaries (dashed lines)(Tetley et al., 2016). After completion of GBE, planar polarisation is lost except at PSBs (thick green lines, stages 9-10), where it is maintained throughout extended germband stages.

Next, we used *in situ* Hybridisation Chain Reaction (HCR) (Choi et al., 2018) to characterize the expression patterns of the 31 candidate genes during early embryogenesis relative to the parasegmental boundaries. HCR has several advantages over traditional *in situ* hybridization: i) because there is no enzymatic amplification, the signal is tightly localized within expressing cells, which helped identify the boundaries of expression with precision, ii) it enables the expression of several genes to be examined simultaneously and iii) it can be combined easily with antibody staining, which we used here to label the cell membranes to facilitate gene expression mapping relative to boundaries. We focused on the period of embryogenesis from stage 5 (late cellularisation) to stage 10 (extended germband) (Figure 1C). From 31 genes, we confirmed that 19 genes were expressed in AP stripes (Sup. Fig. 1), whereas 12 were not (Sup. Fig 2) and were excluded from the candidate list. The 19 genes recovered included the three genes encoding the Toll-like receptors, Toll-2, Toll-6 and Toll-8, already identified by (Pare et al., 2014), thus validating our screening approach. The strength and type of stripe patterns varied between the genes. Some were expressed in seven clear stripes at stages 5 to 7, indicating that they are likely under pair-rule gene network regulation: *ama, best1, comm, comm2, impl2, sca, toll-2, toll-6, toll-8* and *tartan* (Figure 1C and Sup. Fig. 1). Some genes also show clear expression in every parasegment at later stages 9-10, when the germband is extended, either doubling their periodicity from an initial expression in seven stripes (*comm*, *comm2*, *impl2*, *toll-2*) or initiating expression in every parasegment (*dnt, drl, sli*) (Sup. Fig. 1). For the genes with the clearest stripy expression, it was possible to check the position of the stripes relative to the parasegmental boundary markers *ftz* and *slp1*. We found that *best1, blot, comm, comm2, dnt, impl2, toll-8* and *tartan* mRNA expression borders the parasegment boundaries at some point between stages 5 and 10 (Sup. Fig. 3). We decided to focus further investigations on *comm, comm2* and *tartan*, as these genes were most clearly bordering the parasegmental boundaries and were also strongly expressed (Figure 1C).

### A requirement for Tartan in actomyosin enrichment and boundary straightness at PSBs during germband extension

Examining *tartan* expression patterns relative to parasegmental boundaries (PSBs) reveals that, from all the 19 candidates, *tartan* is the gene matching our initial prediction the closest (Tetley et al., 2016)(Figure 2). Like the Toll-like genes, *tartan* encodes a LRR receptor that localizes at the plasma membrane (Chang et al., 1993). The protein localization pattern visualised by antibody labelling matches the mRNA expression patterns, validating the use of HCR to map boundaries (Sup. Fig. 4A). From stage 5 to stage 7, *tartan* is expressed throughout the even-numbered parasegments and borders even-numbered PSBs at the anterior and odd-numbered PSBs at the posterior of its domains (Figure 2A,B and Sup. Fig. 3). Therefore, *tartan* fulfils our prediction of a single receptor expressed in either even or odd parasegments and bordering every PSB. *tartan*’s expression however is not completely uniform across the even-numbered parasegments, being weaker towards the posterior of each stripe at stage 5 to 7. Towards the end of germband extension (stage 8), *tartan*’s posterior border of expression retracts away from the odd-numbered PSBs (Sup. Fig. 3). This is similar to the dynamics of expression of the pair-rule gene *ftz*, which is known to activate *tartan* expression (Chang et al., 1993). From stage 9 onwards, *tartan*’s uniform expression breaks down along the DV axis and the PSBs are bordered only intermittently (Sup. Fig. 1).

**Figure 2:**
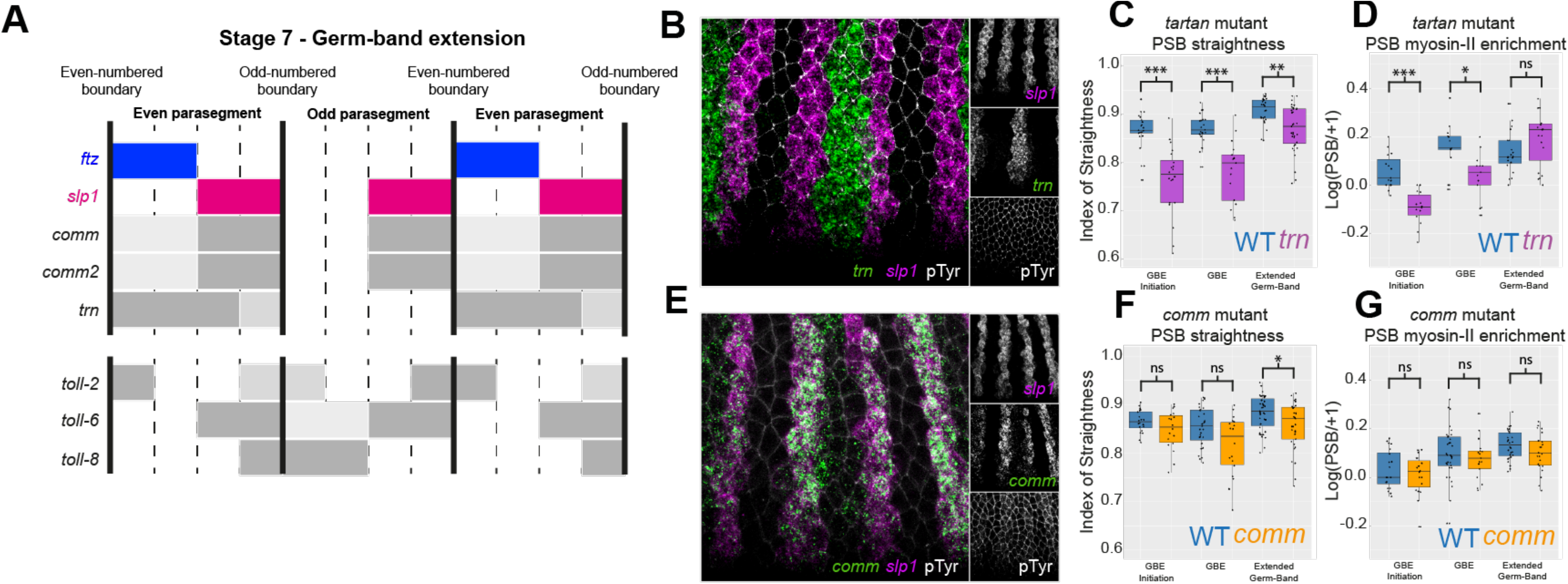
Tartan is required for specifying contractile interfaces at PSB during GBE. A) Diagram based on in situ HCR images (in supplementary figures) showing the expression pattern of *comm, comm2, tartan, toll-2, toll-6* and *toll-8* at the beginning of GBE. Note that the number of cells per parasegmental width varies, with an average of 3.7 cells (Tetley et al., 2016). 4 cells are represented here and subsequent diagrams for simplicity. Dark grey indicate high expression and light grey, lower expression. B) *In situ* HCR of *tartan* expression during GBE. *tartan* is expressed within each even-numbered parasegments and abuts both odd and even-numbered PSBs. C) Index of straightness measures in wildtype (WT)(n= 100 PSBs, from 28 embryos) and *tartan* mutant embryos (n=76 PSBs, from 24 embryos). The convention for P values for this graph and all subsequent graphs are: NS: p>0.05; *p<0.05; **p<0.01; ***p<0.001. D) Myosin II enrichment measures at PSBs in WT (n=96 PSBs and 27 embryos) and *tartan* mutant embryos (n=64 PSBs and 21 embryos). E) *In situ* HCR of *comm* expression during GBE. *comm* is expressed in a similar pattern to *sloppy-paired* during GBE and abuts both odd and even-numbered PSBs. F) Index of straightness measures in WT (n=107 PSBs, from 33 embryos) and *comm* mutant embryos (n=109 PSBs for 30 embryos). G) Myosin-II enrichment measures at PSBs in WT (n=96 PSBs, from 31 embryos) and *comm* mutant embryos (n=91 PSBs, from 28 embryos).

Next, we quantified both Myosin II enrichment (using Sqh-GFP, a knock-in reporter for Myosin II Regulatory Light Chain, (Proag et al., 2019)) and boundary straightness in a *tartan* null mutant (*trn^28.4^*), as a time course from stage 6 to 11 (Figure 2C,D)(Methods). The analysis shows a clear requirement for *tartan* for Myosin II enrichment at PSBs during germband extension, this requirement being strong at GBE onset and diminishing, with no contribution during extended germband stages (Fig. 2D). PSB Straightness quantifications followed a similar trend (Fig. 2C). This matches remarkably the dynamics of expression of *tartan* mapped by HCR, suggesting that the specification of contractile cell-cell interfaces results from an immediate read-out of Tartan receptor asymmetries (Sup. Fig. 1 and 3).

*comm* and *comm2* are duplicated genes located next to each other in the genome and exhibit identical expression patterns, which are markedly different from those of *tartan* (Figure 1C, Sup. Fig. 1 and 3). At stage 5 and 6, their expression in seven stripes is not bordering any PSB and instead straddles the even-numbered PSBs. From stage 7 onwards, *comm* and *comm2* expression doubles in periodicity and the expression becomes localized to the second half of every parasegment, matching *slp1* expression to border the anterior side of each PSB. *comm* encodes a short transmembrane protein that does not localize to the cell surface but regulates the cell surface localization of the receptor Robo and possibly other receptors (Ing et al., 2007; Keleman et al., 2002). Comm protein localizes in puncta consistent with its known ER/Golgi localization, and form a strippy pattern that matches the RNA expression (Sup. Fig. 4B). Comm2 has not been characterized but its amino-acid sequence presents homology with Comm in key domains (Justice et al., 2017). We quantified both Myosin II enrichment and boundary straightness in a *comm* null mutant (*comm^E39^*) (Figure 2F,G). We did not find a significant difference in Myosin II enrichment at any stage, but cannot rule out that an effect of Comm might be masked by a redundancy with Comm2.

### Toll-2 is regulated by Wingless signalling and required for Myosin II enrichment and boundary straightness at PSBs during extended germband stages

In addition to looking at germband extension, we also wanted to assess PSB function at extended germband stages (stages 9-11). At these stages, actomyosin enrichments initiated during GBE at PSBs are maintained by Wingless signalling (Monier et al., 2010). In *wingless* mutants, both actomyosin enrichment and boundary straightness are lost at PSBs, as is the elevated tension along the cell-cell interfaces, shown by laser ablation experiments (Monier et al., 2010; Scarpa et al., 2018; Urbano et al., 2018). We reasoned that any cell surface receptor contributing to maintaining actomyosin enrichments at PSBs must be under Wingless signalling regulation. We therefore performed HCR in a *wingless* null mutant (*wg^CX4^*), for the candidate genes expressed in stripes at extended germband stages (*best1, comm, comm2, dnt, drl, sli*) as well as for *tartan* and the *toll-like* receptors (Sup. Fig. 5). Of these 11 genes, only *toll-2* lost expression in *wg^CX4^ embryos* compared to wildtype (Figure 3A,B and Sup. Fig. 5). To confirm regulation by Wingless signalling, we examined *toll-2* expression (alongside the other *toll-like* receptors and *tartan* as controls) in embryos ubiquitously expressing Wingless (*armGal4/UASwg*). Again, *toll-2* was the only gene responding robustly to Wingless signalling, its expression broadening towards the posterior to reach the anteriorly broadened expression of the *slp* domain (Figure 3C). The broadening of *toll-2* expression matches the known changes in *engrailed* expression in *armGal4/UASwg* embryos (Scarpa et al., 2018; Urbano et al., 2018), confirming that Wg signalling regulates *toll-2* expression. The *toll-2* expression during GBE is known to straddle the PSBs (Figure 2A) (Pare et al., 2019; Pare et al., 2014). This is the case also at extended germband stages (Figure 3A). We quantified both Myosin II enrichment and boundary straightness in a *toll-2* null mutant (Figure 3D,E). As in (Lavalou et al., 2021; Pare et al., 2019; Pare et al., 2014), we do not detect a contribution for *toll-2* during germband extension, however, we do find a contribution of *toll-2* at extended germband stages. This suggests that the role of Wingless signalling in maintaining contractile interfaces at PSBs is mediated, at least in part, by Toll-2.

**Figure 3:**
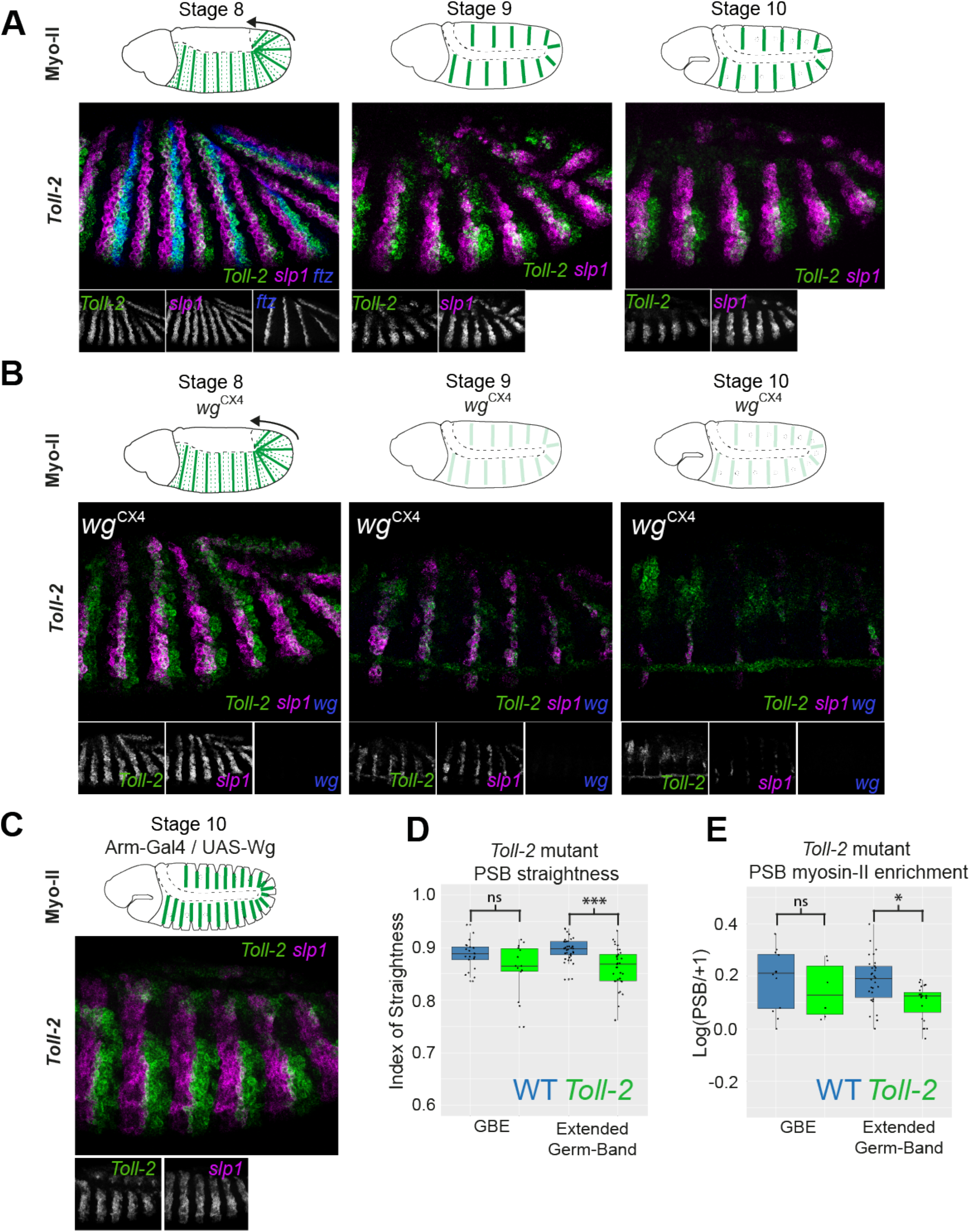
Toll-2 is required for PSB straightness at germband extended stages of *Drosophila* embryogenesis. A) *In situ* HCR of *toll-2* expression in WT embryos showing that *toll-2* is expressed in AP stripes immediately posterior to *slp* expression (which labels the anterior edge of PSBs) during GBE (stage 8) and at extended germband stages (stages 9 and 10). Note that Toll-2 slightly straddles the *slp* expression domain and thus the PSBs (merged signal in white). B) *In situ* HCR of *toll-2* expression in *wingless* mutant embryos. At stage 8, *toll-2* expression is as in WT. At stage 9, *toll-2* expression is markedly reduced and by stage 10, *toll-2* expression is absent. C) *In situ* HCR of *toll-2* expression during GBE in embryos overexpressing *wingless. toll-2* expression responds to Wingless signalling by expanding posteriorly, filling about half of each parasegment. **D)** Index of straightness measures in WT (n=103 PSBs, from 24 embryos) and *toll-2* mutant embryos (n=109 PSBs, from 26 embryos). E) Myosin II enrichment measures at PSBs in WT (n= 86 PSBs, from 24 embryos) and *toll-2* mutant embryos (n=89 PSBs, from 24 embryos).

We conclude that our screening approach has allowed us to identify a contribution of *tartan* and *toll-2* to the PSB’s mechanical properties during early embryogenesis. They have distinct windows of requirement, with *tartan* contributing during germband extension and *toll-2* contributing during extended germband stages under Wingless signalling control.

### Tools to monitor boundary activity during germband extension in live embryos

Our analysis in fixed embryos suggests that Tartan’s requirement is limited to convergent extension, matching closely its window of expression at PSBs. This is intriguing because it suggests that the specification of contractile interfaces is a rapid and short-lived response to Tartan’s asymmetric expression at boundaries. In order to analyse more precisely the dynamics of requirement for *tartan*, we developed tools to monitor boundary mechanical properties during axis extension. In previous work, we had quantified Myosin II polarity in live embryos using Sqh-GFP (the reporter for Myosin II Regulatory Light Chain), while tracking cells with the cell membrane marker Gap43-Cherry (Tetley et al., 2016). Here we develop boundary straightness measurements in real time as a functional assay for boundary mechanical properties and a proxy for actomyosin enrichment.

To follow the dynamics of boundary straightness in live embryos, we took advantage of the MS2-MCP system implemented in *Drosophila* embryos (Garcia et al., 2013), to label in real time the transcription of the parasegmental boundary marker *engrailed* (Sup. Fig. 6A). The reason for using a transcriptional read-out rather than a protein reporter is that we found that tagged proteins of segmental markers, constructed by others or ourselves, do not give a fluorescent signal strong enough for tracking parasegments in live embryos. One exception is *eve-YFP*, which we used previously, but has the limitation of labeling only alternate parasegmental boundaries (Tetley et al., 2016). We fused a 2099bp region upstream of the *engrailed* promoter, the VT15159 enhancer, to a MS2 reporter containing 24 MS2 stem loops and *lacZ*, generating the construct EnVT15159-MS2 (Sup. Fig. 6B and Methods). We checked by HCR that *lacZ* expression from this construct recapitulates the endogenous pattern of *engrailed* expression during axis extension (Sup. Fig. 6C,D). The only differences are brighter Ftz-positive stripes (marking even-numbered parasegments), which might be a consequence of the known delay in transcription initiation of *engrailed* in odd-numbered parasegments compared to even-numbered ones (DiNardo et al., 1988).

Next, we visualised transcription from EnVT15159-MS2 by co-expressing MCP-GFP, which binds to the 24 MS2 stem loops in nascent transcripts and give rise to fluorescent dots corresponding to *engrailed* transcription in the nuclei of live embryos (Figure 4B and Sup. Fig. 6A). A kymograph of the dots reveals that as for *lacZ* expression from the same reporter (Sup. Fig 6C,D), reported *engrailed* transcription is brighter in alternate parasegments (Figure 4B’). This is useful as it gives us a mean to distinguish even-numbered from odd-numbered parasegments. To associate *engrailed* transcriptional dots with given cells, we used Gap43-mCherry as previously to label cell membranes and track cell positions (Figure 4C)(Tetley et al., 2016). We developed additional computational methods to track the transcriptional dots in order to identify *engrailed* positive cells (Figure 4C’) and thereby the cell-cell interfaces of the parasegmental boundaries (Sup. Fig. 7 and Methods). As shown below, we found that the parasegmental boundaries identified by these methods are, as expected, significantly straighter than control (non-boundary) interfaces throughout axis extension, thus validating the use of a reporter of *engrailed* transcription to identify parasegmental boundaries in live embryos.

**Figure 4:**
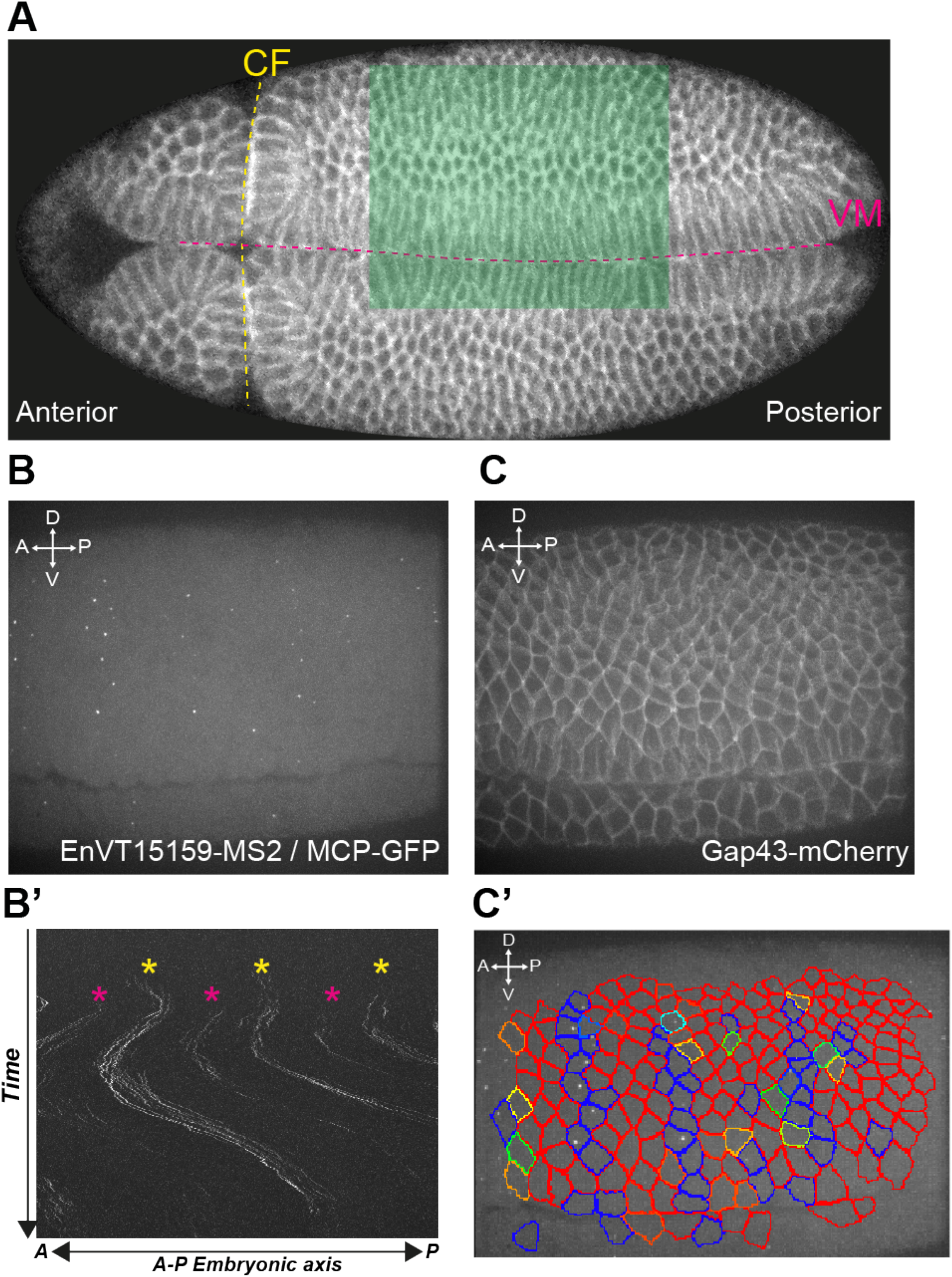
Tracking parasegmental boundaries in live *Drosophila* embryos. A) Spinning disc confocal image of a live gastrulating *Drosophila* embryo expressing Gap43-mCherry. The cephalic furrow (CF) and ventral midline (VM) are annotated with dashed colored lines. The ventrolateral region shaded green is the approximate area imaged via spinning disc confocal microscopy. (Anterior = Left, Posterior = Right. 20x magnification). B,C) Maximum intensity projection of a single time frame from an embryo expressing the transgenes EnVT15159-MS2, MCP-GFP and Gap43-mCherry. MCP-GFP binds the MS2 loops on nascent mRNA transcribed from the *engrailed* enhancer EnVT15159 and form spatially localized fluorescence within nuclei that resemble dots. B’) Kymograph showing the distribution of transcriptional dots along the embryonic axis during GBE. Magenta asterix indicates stripes that coincide with odd-numbered PSBs, while yellow asterix indicates the brighter stripes that coincide with even-numbered PSBs. C) Gap43-mCherry protein localizes to cell membranes. C’) Movie frame showing cell segmentation based on the Gap43-mCherry signal and boundary cell tracking based on the MCP-GFP dot signal. Assigning the dots to cells reveals the position of Engrailed-expressing cells.

### Tartan is required for parasegmental boundary mechanical activity throughout axis extension

To compare parasegmental boundary straightness, we analysed 3 movies each of wild-type and *tartan* mutant embryos carrying the transgenes *EnVT15159-MS2, MCP-GFP* and Gap43-mCherry. Movies were acquired as before (Tetley et al., 2016)(see field of view in Figure 4A), cells were segmented automatically based on the Gap43-mCherry signal, then segmentation was corrected manually (Methods). Manual correction was important to recover enough cell-cell interfaces for the boundary straightness analysis. *engrailed* transcriptional dots from *EnVT15159-MS2/MCP-GFP* were tracked to locate the *engrailed* stripes and to find the parasegmental boundaries at the anterior border of each stripe (Sup. Fig. 7). We then compared the angle of cell-cell interfaces relative to the antero-posterior (AP) axis for parasegmental boundary interfaces and for control interfaces located one cell diameter posteriorly or anteriorly (+1 and −1 interfaces, respectively) (Figure 5A-C). In wild-type embryos, cell-cell interfaces are quite straight at the beginning of axis extension, with 55 to 65% of PSB and control interfaces having an angle greater than 60 degrees relative to the AP axis (Figure 5D). This initial interface straightness is caused by the invaginating mesoderm pulling on the ventral border of the ectoderm around the time axis extension starts (Butler et al., 2009; Lye et al., 2015). Once the mesoderm has invaginated, the tissue relaxes and the control interfaces lose their alignment for the remainder of axis extension. In contrast, parasegmental boundaries remain aligned throughout axis extension, with 60% of boundary interfaces having an angle greater than 60 degrees relative to the AP axis (Figure 5D,D’). These trends are remarkably similar to our previous measurements for even-numbered PSBs identified using Eve-YFP (see fig2 K, L in (Tetley et al., 2016), validating our new method to identify boundaries, and confirming that parasegmental boundaries behave as mechanical boundaries during *Drosophila* germband extension.

**Figure 5:**
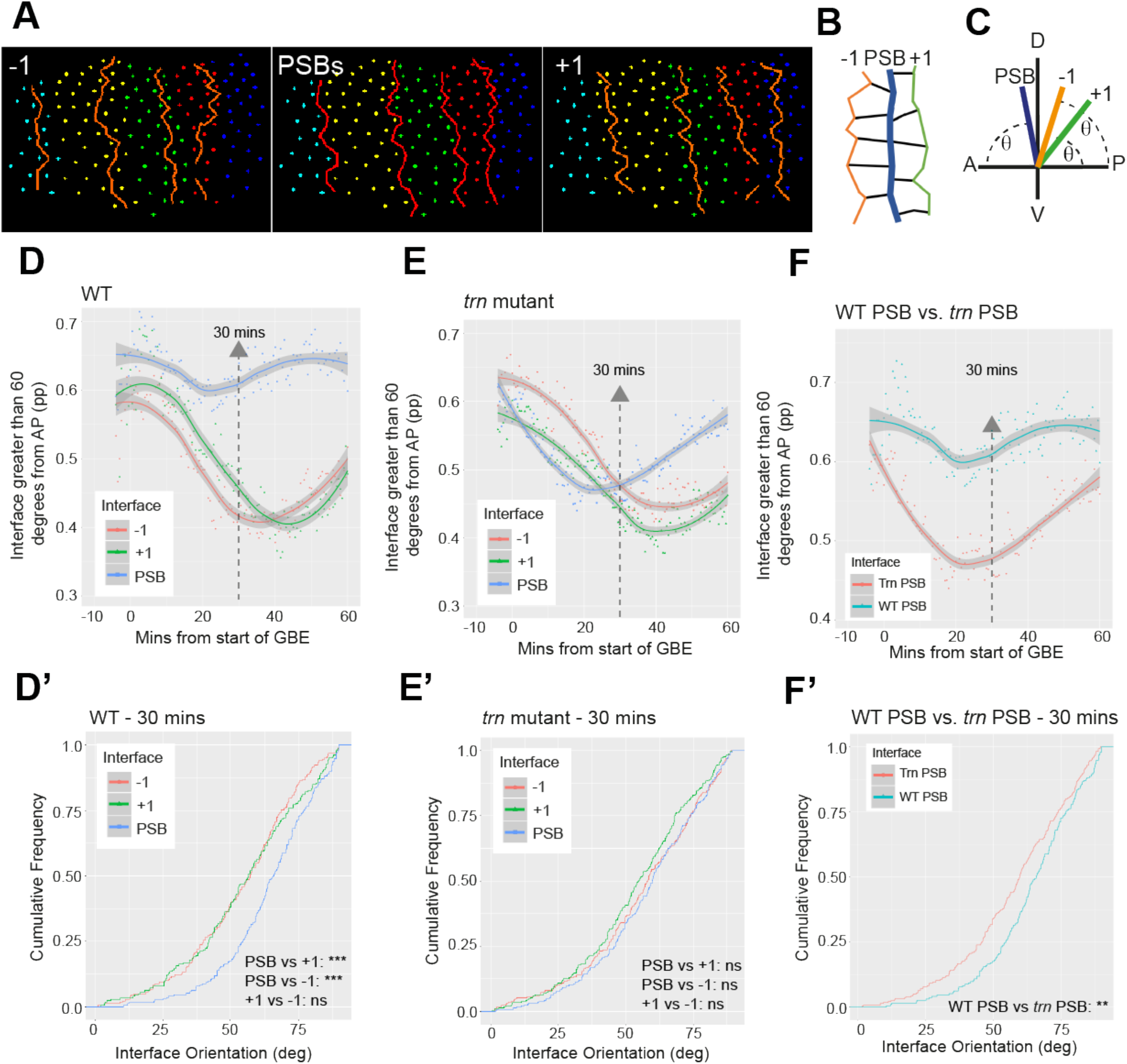
Live imaging reveals Tartan is required for PSB straightness throughout germ-band extension. A) In tracked movies, cell centroids colored by parasegment identity. PSBs, −1 and +1 interfaces are overlayed to show their position relative to one another. B) Diagram showing the relative position of −1, PSB, and +1 interfaces. −1 and +1 interfaces correspond to the column of AP interfaces one cell diameter to the anterior and to the posterior of PSBs, respectively. C) Diagram showing how the orientation of cell interfaces that make up each column is calculated. Theta represents the measured angle between the embryonic AP axis and each cell interface. Angles of each cell interface that make up a column are measured automatically in our tracking software. D, E) Plots showing the proportion of interfaces at PSB and control −1 and + 1 interfaces that are greater than 60 degrees from the AP axis, in the course of GBE, for 3 wildtype (D) and 3 *tartan* mutant embryos (E). A loess curve (span 0.75) has been fitted to the data. D’, E’) Statistical comparison at time point 30 minutes with a Kolmogorov-Smirnov non-parametric test undertaken on the cumulative frequencies of interface angles. F) Subset of D and E curves to directly compare the PSB straightness in WT and *tartan* mutant embryos in the course of GBE, with statistics for timepoint 30 minutes in F’.

In *tartan* mutants, the control interfaces anterior and posterior to the PSBs (−1 and +1) show the same behaviour as in wild-type embryos: they are initially aligned at the start of germband extension by the mesoderm pull, then lose their alignment once the tissue relaxes. Strikingly however, the straightness of parasegmental boundaries in *tartan* mutants is indistinguishable from control interfaces 30 minutes into axis extension (Figure 5 E,E’). Direct comparison between *tartan* and wildtype PSBs demonstrate that PSBs in *tartan* null mutants have lost their straightness throughout axis extension (Figure 5F,F’). These measurements demonstrate that *tartan* is required for PSB straightness, and thus actomyosin enrichment at these boundaries, during axis extension. The timings of the loss of straightness is consistent with our time-course of actomyosin enrichment and boundary straightness in fixed embryos (Figure 2C,D). Note that at the beginning of axis extension, PSBs in *tartan* mutants lose their alignment more quickly than control interfaces, consistent with a loss of actomyosin enrichment at PSBs in *tartan* mutants from early on (Figure 5E). After 40 minutes into germband extension, *tartan* PSB straightness curves start to increase towards wildtype, which is again consistent with fixed embryos time-courses (Figure 2C,D) and suggest that other receptor systems take over then to promote actomyosin enrichment at PSBs. It also suggests that boundary straightness is an immediate read-out of the molecular asymmetries present at a given period of development.

### Pair-rule regulation of *tartan* during germband extension

*tartan* is known to be under pair-rule regulation, in particular from Ftz (Chang et al., 1993). In a *ftz* mutant, *tartan* stripy expression along AP is lost at gastrulation and we confirm this by HCR (Sup. Fig. 8). This leaves us with a conundrum however: our results suggest that Tartan is required for the mechanical activity of every PSB, while only even-numbered PSBs are known to be lost in *ftz* mutants (Larsen et al., 2008). We first checked whether *tartan* is indeed required at all PSBs, by classifying odd and even-numbered PSBs using the stronger signal for *engrailed* transcriptional dots in even-numbered PSBs. We find that the loss of straightness at PSBs in *tartan* mutants is the same at odd versus even-numbered PSBs (Sup. Fig. 9A,A’), confirming that *tartan* is required for actomyosin enrichment at all parasegmental boundaries during axis extension. We also looked at the straightness of both classes of PSBs in wildtype: even-numbered PSBs are slightly straighter than odd ones throughout extension, which might reflect the fact that *tartan* expression abuts the even-numbered PSBs more closely than the odd ones, so a Tartan receptor asymmetry might be more pronounced at even-numbered PSB (Sup. Fig 9B,B’). Our statistical tests however do not detect a significant difference, so we conclude that the enrichment of actomyosin is roughly similar at every PSB during axis extension and that *tartan* is required for this enrichment at all PSBs.

Next, we analysed boundary straightness in *ftz* RNAi knockdown embryos, using the same methods as for *tartan* mutants. *ftz* dsRNA injection resulted in pair-rule cuticle phenotypes identical to those of *ftz* null mutants, showing that we have an efficient knockdown (Sup. Fig. 9D,E). We analysed 3 movies each of *ftz* dsRNA injected and control buffer injected embryos. *engrailed* transcriptional dots in buffer injected embryos have the same pattern as in wildtype, with brighter even-numbered stripes of *engrailed* dots (Figure 6 A,C). Straightness curves are similar to wildtype, with PSBs consistently straighter than control interfaces throughout axis extension (Figure 6E, E’). In the *ftz* knockdown, we expect the even-numbered stripes to be lost, since Ftz is required to activate *engrailed* transcription in even-numbered parasegments (Florence et al., 1997; Howard and Ingham, 1986). Consistent with this, we find that alternate stripes of *engrailed* transcriptional dots are gone, with weak stripes remaining, which we infer are the odd-numbered stripes (Figure 6B). Although weak, we were able to use those traces to track the odd-numbered PSBs in *ftz* knockdown embryos (Figure 6D). These odd-numbered PSBs are clearly straighter than control interfaces (Figure 6F,F’). Next, we compared directly odd-numbered PSB straightness in buffer-injected versus *ftz* RNAi embryos (Figure 6G,G’). We find that both have the same straightness, indicating that as expected from the known role of Ftz in parasegment patterning, odd-numbered PSBs are behaving as in wildtype.

**Figure 6:**
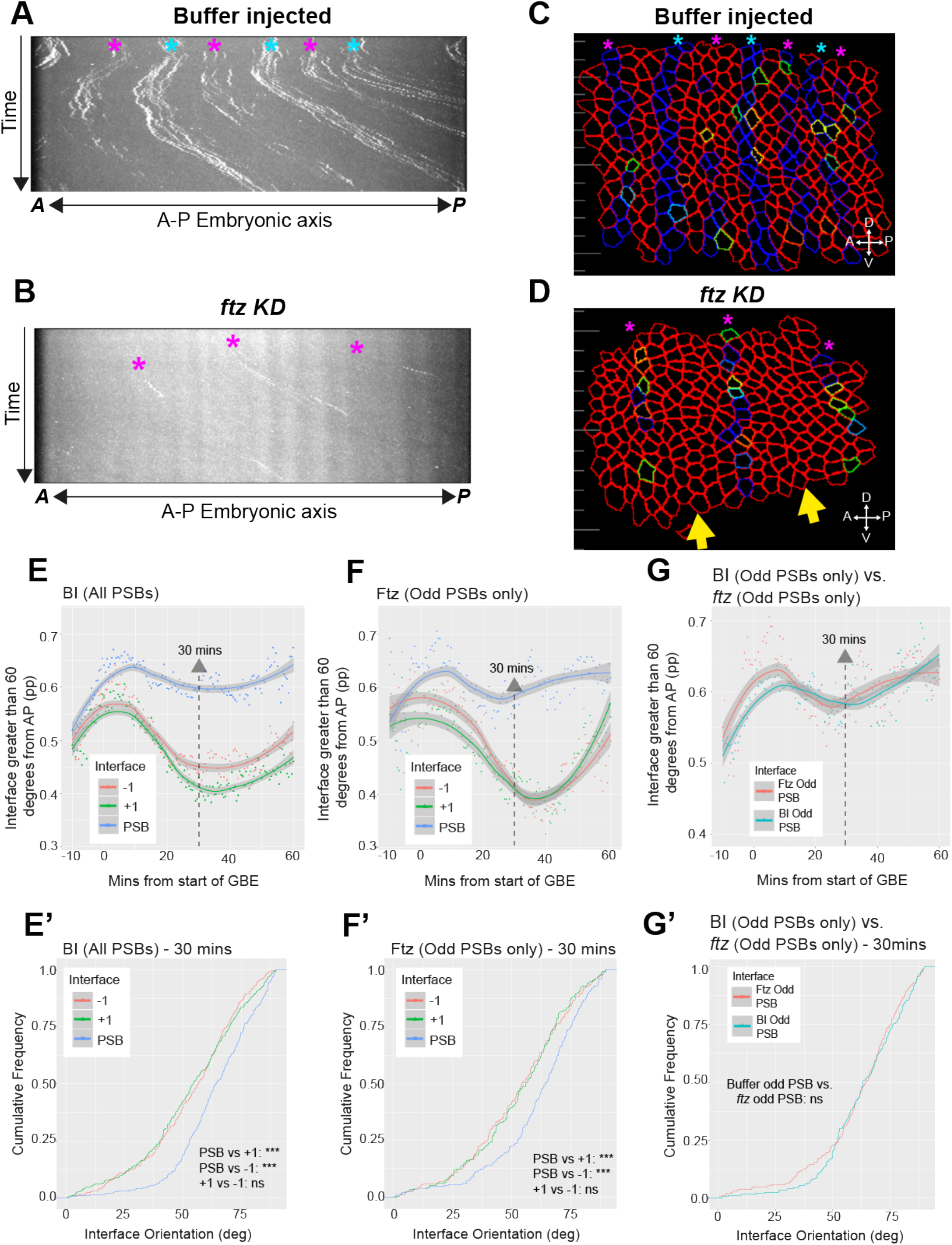
Boundary tracking in live embryos reveals that odd-numbered PSBs in *ftz* mutant embryos retain their straightness during GBE. A,B) Kymographs showing the spatiotemporal distribution of transcriptional MCP-GFP dots marking the PSBs during GBE. Magenta asterix indicates stripes that coincide with odd-numbered PSBs, while blue asterix indicates the brighter stripes that coincide with even-numbered PSBs. In A, buffer injected embryos show the same pattern as wildtype embryos (compare with Fig. 4B’), while in B, embryos injected with *ftz* dsRNA (*ftz* KD) have, as expected, lost the stronger stripes that coincide with even-numbered PSBs. C,D) Movie frames showing examples of boundary tracking in a buffer injected embryo (C) and a *ftz* KD embryo (D). Yellow arrows indicate the approximate position where even-numbered *engrailed* positive stripes are missing in *ftz* KB embryos. E, F) Straightness measurements in the course of GBE for 4 buffer injected (BL) and 4 *ftz* KD embryos. Plots show the proportion of interfaces that are greater than 60 degrees from the AP axis, for PSBs and +1 and −1 control interfaces. A loess curve (span 0.75) has been fitted to the data. Note that for *ftz* KD (F), only the odd-numbered PSBs and corresponding controls interfaces could be tracked. E’, F’) Statistical comparison for timepoint 30 minutes for plots E and F uses a Kolmogorov-Smirnov non-parametric test undertaken on the cumulative frequencies of interface angles. G) Subset of data from E and F curves to directly compare the straightness of odd-numbered PSBs in buffer injected and *ftz* KD mutant embryos in the course of GBE, with statistics for timepoint 30 minutes in G’.

The above analysis of *tartan* and *ftz* mutants confirm that we have a paradox: all parasegmental boundaries are affected in *tartan* mutants, whereas in *ftz* mutants, the odd-numbered PSBs have the same mechanical properties as in wildtype. However, in *ftz* mutants, the AP stripes of *tartan* expression are gone (note that there is residual expression in the DV axis, Sup. Fig 8). One possible explanation is that in a *ftz* mutant, changes in pair-rule gene expression changes the expression of cell surface receptors, which in turn rescues the boundary mechanical activity and thus actomyosin enrichment at odd-numbered PSBs. We find that the expression of all three Toll-like receptors *toll-2, toll-6* and *toll-8* is changed in *ftz* mutants (Sup. Fig. 8). Registration with *wg* expression suggests that *toll-2* and *toll-6* are expressed coincidently with the remaining Wg stripes in odd-numbered parasegments, while *toll-8* is expressed in a complementary pattern (Sup. Fig. 8). This altered expression provides the potential for any of the three Toll-like receptors, individually or in combination, to rescue the loss of Tartan at odd-numbered PSBs in *ftz* mutants.

## Discussion

Our systematic approach to find genes required for parasegmental boundary formation in *Drosophila* embryos identifies Tartan and Toll-2 as cell surface receptors key for mechanical boundary activity at distinct developmental times: Tartan is required under pair-rule gene control during germband extension and Toll-2 is required under Wingless signalling control during the extended germband stages.

One of our motivations for taking a systematic approach was to evaluate how many cell surface receptors are required for compartmental boundary formation during axis extension. In (Tetley et al., 2016), we proposed that a single receptor expressed in either even- or odd-numbered stripes would be the minimal number. Tartan fits this single receptor hypothesis because it is expressed in even-numbered parasegments and is required for polarized contractility of interfaces at every parasegment boundary during axis extension (this study and (Pare et al., 2019). We have identified six other genes that encode cell surface receptors or regulators of cell surface receptors and that are asymmetrically expressed at parasegment boundaries during axis extension: *Best1, blot, comm, comm2, dnt* and *ImpL2*. The removal of *comm* on its own did not show a significant contribution to interfacial contractility at PSBs (Figure 2F, G). *toll-6* and *toll-8* are also expressed differentially at PSBs, at least some of the time (Sup Fig. 3), but previous reports showed that these do not contribute significantly to interfacial contractility at PSBs during germband extension (Lavalou et al., 2021; Pare et al., 2019). Although we cannot rule out a contribution of the receptors that have not been tested yet, based on the robust phenotype found in tartan mutants, it is likely that if any, this contribution would be minor at PSBs during GBE.

Indeed, our analyses in both fixed and live embryos show that Tartan is required for PSB actomyosin enrichment from the very start of GBE. In live embryos, PSB straightness in *tartan* mutants is in fact lesser than flanking columns of interfaces during the initiation of GBE (stage 7, until approximately 20-25mins in GBE, Figure 5E). This is consistent with our Myosin II quantifications at stage 7 in fixed *tartan* mutant embryos, which shows that Myosin II is depleted at PSBs relative to the interfaces one diameter posteriorly (+1 interfaces) (Figure 2D, left data set, pink box). In contrast, in wild-type embryos, PSBs and +1 interfaces have the same relative enrichment (Figure 2D, left data set, blue box), which is consistent with our quantitative analysis of Myosin II polarization during GBE, where all cells initially display a bipolarity in Myosin II (Tetley et al., 2016). Planar polarization at AP cell-cell interfaces other than PSBs requires the Toll-like receptors *toll-2*, *toll-6* and *toll-8* (Lavalou et al., 2021; Pare et al., 2019; Pare et al., 2014). So our analysis of PSBs and interfaces one cell diameter away (−1 or +1 flanking interfaces) suggests that in *tartan* mutants PSBs interfaces do not enrich Myosin II at the start of GBE, whereas the other AP interfaces enrich Myosin II normally. Together, these findings are consistent with *tartan* encoding the key molecular asymmetry required for PSB contractility during axis extension.

Although actomyosin enrichment at PSBs is initially indistinguishable from other AP interfaces at the beginning of GBE, it becomes more prominent as germband extension progresses (Tetley et al., 2016) (Figure 1C). This change is detected in our Myosin II quantifications in fixed wild-type embryos in this study: as mentioned above, there is no relative enrichment between PSBs and +1 interfaces at the beginning of GBE (Figure 2D,G; left data sets, blue boxes). In contrast, mid-GBE, we can systematically detect an enrichment at PSBs relative to +1 interfaces (Figure 2D, G; data sets in center, blue boxes; also Figure 3E, left data set, blue box). Our interpretation for the increase in PSB/+1 relative enrichment is that as cells intercalate in the germband, the initial bipolarity in Myosin II changes to unipolarity, with only one side of each cell (either anterior or posterior) remaining enriched in Myosin II (Tetley et al., 2016). We have argued that this supports a model where cell-cell interactions between cells of different “identities” along AP generate planar polarity, because once cells have intercalated, pairs of cells with the same initial AP positional information do not appear to re-enrich Myosin II significantly at their new shared interface. So based on these earlier findings, we expect the PSB/+1 relative enrichment to increase as germband extension progresses, as we find here.

In our previous study, once the germband cells had intercalated significantly, we could detect Myosin II enrichment every two cell diameters, identifying two intra-parasegmental boundaries (Figure 1C)(Tetley et al., 2016). Interfaces located at intra-parasegmental boundaries enrich Myosin II more and are straighter than their respective +1/-1 interfaces, so they behave as mechanical boundaries. Interestingly, these two intra-parasegmental boundaries have comparable Myosin II enrichment and straightness, but those are significantly lower compared to PSBs (Tetley et al., 2016). The mechanisms causing stronger Myosin II enrichment at PSBs compared to intra-parasegmental boundaries are unknown. Taking together findings from (Lavalou et al., 2021; Pare et al., 2019; Pare et al., 2014) and this study, we can infer that these three mechanical boundaries are generated by distinct sets of receptor asymmetries: the PSB by Tartan asymmetries, and the two intra-parasegmental boundaries by Toll-like 2,6,8 asymmetries. One possibility is that there is some undetected contribution of Toll-like asymmetries at PSBs and that Tartan is added “on top”, explaining the stronger actomyosin enrichment at PSBs. Alternatively, the signalling in response to Tartan asymmetry might be more effective at increasing actomyosin contractility. Recent reports have identified distinct receptor interactions for Tartan and the Toll-like receptors, which might suggest distinct signalling pathways. Tartan interacts with the teneurin Ten-M (Pare et al., 2019), while Toll-8, and likely Toll-2, interacts with the adhesion GPCR Cirl (Lavalou et al., 2021). Signalling pathways downstream Tartan/TenM interactions are unknown, but recently Src and Pi3K signalling have been identified downstream Toll-like 2 asymmetries (Tamada et al., 2021). Signals downstream Tartan and Toll-like receptors must eventually regulate actomyosin contractility, which is known to be controlled in a quantitative manner by specific GPCR receptors in the germband (Garcia De Las Bayonas et al., 2019; Kerridge et al., 2016).

Whatever the mechanisms of actomyosin enrichment, our study shows that Tartan is not the only receptor able to trigger interfacial contractility at PSBs. Rather, our findings suggest that receptors might be interchangeable, and it is their expression patterns that determine at which cell-cell interfaces receptors are required for interfacial contractility. First, we find that odd-numbered PSBs remain contractile and form mechanical bounadries in *ftz* mutants, despite the loss of *tartan*’s AP stripe pattern in these mutants. In *ftz* mutants, the pattern of every Toll-like receptor changes, and it is therefore possible that one or a combination of these receptors rescues interfacial contractility at odd-numbered PSBs. In future, this could be addressed by removing candidate Toll-like receptors in *ftz* mutant to search which combination abolishes interface contractility at odd-numbered PSBs, although we cannot exclude that another, unknown receptor takes over. Second, in *tartan* mutants, the PSBs progressively recover their straightness towards the end of GBE. This is coincident with a breakdown in *tartan* expression along PSB borders and suggests that other inputs take over from Tartan at extended germband stages. A good candidate for this later input is Toll-2, which we have identified as being required for PSBs actomyosin enrichment at extended germband stages. We find that Toll-2 expression during this period of development is under Wingless signalling regulation. Since actomyosin enrichment maintenance at PSBs during extended germband stages requires Wingless signalling (Monier et al., 2010; Scarpa et al., 2018; Urbano et al., 2018), this suggests that Toll-2 mediates this role. Thus a minimal hypothesis can be constructed where contractile interfaces at PSBs are first specified during axis extension by Tartan receptor asymmetries controlled by pair-rule gene regulation (mainly *ftz*). Then as Wingless signalling becomes active, Toll-2 takes over from Tartan to maintain contractility at PSB boundary interfaces. Our live imaging analysis also suggests that mechanical boundary formation responds in real time and with high sensitivity to molecular asymmetries, because the recovery of PSB straightness in *tartan* mutant parallels the loss of expression of *tartan* along PSBs. It is likely that mechanosensitive feedbacks contribute to this responsiveness; indeed, Myosin II-enriched cell interfaces connected to each other enrich more Myosin II and are under greater tension than isolated interfaces, both during germband extension (Fernandez-Gonzalez et al., 2009) and extended germband stages (Scarpa et al., 2018), suggesting the existence of a positive mechanosensitive feedback. Consistent with this notion, in both cases, decreasing tension at connected cell-cell interfaces using laser cuts also decreases Myosin II enrichment (Fernandez-Gonzalez et al., 2009; Scarpa et al., 2018). So it is possible that mechanosensitive feedback increase actomyosin enrichment along PSBs, contributing to the real-time responsiveness of boundary formation. This might also contribute to the robustness of boundary formation (Martin et al., 2021).

Our *in silico* screen was based on the assumption that differential expression of receptors underlies interfacial contractility at PSBs. One limitation of our approach is that borders in mRNA expression detected by HCR do not necessarily equate with an asymmetry in protein localization, since post-translational regulation could modulate receptor localisation. However, this approach was sufficient to identify Tartan and Toll-2. Also, comparison of protein and mRNA expression patterns for *tartan* and *comm* suggests that these are comparable, with the main difference being that the mRNA pattern is ahead in time. For example, at the beginning of GBE *tartan* mRNA expression retracts away from the odd-numbered PSBs (as does *ftz*) while the protein pattern is still abutting the PSBs (Sup. Fig. 4). Recent reports show that the asymmetric localization of the LRR receptors at the protein level is essential for the formation of contractile interfaces (Lavalou et al., 2021; Pare et al., 2019). The cell surface interactors Ten-M (for Tartan) and Cirl (for Toll-8) have a uniform RNA expression in embryos and were eliminated as candidates in our *in silico* screen (Sup. Fig. 2). At the protein level, Ten-M and Cirl become localized at boundary cell-cell interfaces via their interactions with Tartan and Toll-8, respectively (Lavalou et al., 2021; Pare et al., 2019). The planar polarization of those heterophilic receptor complexes is thought to underlie the formation of contractile cell-cell interfaces, via pathways which remain to be fully elucidated (Garcia De Las Bayonas et al., 2019; Tamada et al., 2021). We conclude that searching for asymmetric RNA expression of receptors at boundaries is overall a good strategy to find genes required for boundary formation, but it is not exhaustive and also does not inform precisely about the behaviour of the corresponding proteins at boundary cell-cell interfaces. The latter is illustrated by our findings that Toll-2 is required for actomyosin enrichment at PSBs in extended germband stages and that Toll-2 mRNA expression consistently straddles rather than border the PSBs (and this expression is lost in *wingless* mutants). Thus, Toll-2 mRNA expression does not precisely border the PSBs. One possibility is that regulation of Toll-2 at post-translational level creates planar polarities at the PSB similarly to the earlier GBE stages. Whether Wingless signalling contributes directly to a putative planar polarization of Toll-2, in addition to transcriptional regulation, will need to be addressed.

## Acknowledgements

We would like to thank Erik Clark and Matt Benton for advice about *in situ* HCR; Julia Falo-Sanjuan for advice about the MS2-MCP system; Robert Zinzen, Nikolaos Karaiskos and Nick Brown for providing data for the *in silico* screen; Sarah Bray, Guy Tear and Shigeo Hayashi for reagents; Rob White and Erik Clark for critical reading of the manuscript; all present and past members of the Sanson lab for discussion.

## Methods

### *In silico* screen

To identify *Drosophila* genes expressed in anteroposterior (AP) stripes in the early embryo, the BDGP library (http://insitu.fruitfly.org/cgibin/ex/insitu.pl) was filtered using the descriptors “Pair-Rule” and/or “Segmentally Repeated”, the Fly-FISH library (http://fly-fish.ccbr.utoronto.ca/) was filtered using the descriptors “Pair-Rule” and/or “Segment Polarity”, the Enhancer Library (http://enhancers.starklab.org/) was filtered using the descriptors “A-P Stripes” and/or “Pair-Rule”, and the vISH library (https://shiny.mdc-berlin.de/DVEX/) was manually interrogated for the expression pattern of 441 genes predicted to encode transmembrane adhesion protein in *Drosophila* (Hynes and Zhao, 2000). Manual clustering analysis of the vISH library raw data was also performed to identify the top 200 genes expressed in the same cells as those expressing *even-skipped* or *fushi-tarazu* in early embryos (personal correspondence from Nikos Karaiskos and Robert Zinzen, Max Delbrück Center for Molecular Medicine, Berlin). To identify genes encoding proteins that localise to the cell surface, the UniProt data resource (http://www.uniprot.org) was filtered for the descriptors: *annotation:(type:transmem) AND organism:“Drosophila melanogaster (Fruit fly)”* and also annotation:(type:signal) AND organism:“Drosophila melanogaster (Fruit fly)”. To identify genes regulated by the pair-rule gene network, differentially expressed genes resulting from the knock-down of *even-skipped* and *runt* in early *Drosophila* embryos were obtained from (Pare et al., 2014). Further, the BDTNP database (http://bdtnp.lbl.gov/Fly-Net/) was queried to identify genes neighbouring *fushi-tarazu, sloppy paired1, paired* and *runt* DNA binding sites, while the DroID database (http://www.droidb.org/Index.jsp) was used to identify genes neighbouring *even-skipped*, *hairy* and *odd-skipped* DNA binding sites. A custom R script was used to wrangle the downloaded filtered datasets into a standardised dataframe format and identified genes that fulfilled candidate criteria. Initial candidate genes had their raw *in situ* hybridisation images, contained in each library, manually assessed. If a gene was found not to be expressed in AP stripes the gene was excluded. The ModEncode temporal expression data set (Roy et al., 2010) (annotated version kindly provided by Nick Brown, University of Cambridge) was used to exclude genes with less than 5% of their total embryonic expression (0-24hrs AEL) occurring between 0-6hrs AEL. Finally, a manual investigation of protein localisation and described role was undertaken using Flybase and a search in the primary scientific literature.

### Whole mount *in situ* HCR v3.0

Two to five hours old *yw^67^* embryos were collected on apple juice agar plates at 25°C, fixed in 4% formaldehyde/heptane for 20 min, and stored at −20°C in methanol until required. *In situ* HCR v3.0 with split initiator probes were performed as in (Choi et al., 2018). The probes sets were designed by Molecular Instruments to target exons present within every gene isoform. Embryos for whole mount *in-situ* HCR were first post-fixed in 4% formaldehyde, then washed in PBT (PBS with 0.1% Tween-20), then 5XSSCT prior to hybridisation. Embryos were pre-hybridised in warm hybridisation buffer for 30 mins at 37°C. Embryos were incubated in the probe hybridisation solution (0.8pmol of each probe in 200uL) at 37°C overnight. Following overnight incubation, excess probes were removed by washing in wash buffer at 37°C, then in 5XSSCT at RT. The embryos were pre-amplified in buffer then final amplification solution was added (6pmol of each snap cooled fluorescently labelled hairpin added to 50-100μL of amplification buffer). Embryos were incubated in the amplification solution overnight then washed in 5XSSCT. If an antibody immunostain was to follow, embryos were washed in PBS-TX before being blocked in a PBS-TX-BSA solution as standard. Embryos were mounted in Vectashield (Vectorlabs) before imaging.

### Confocal imaging of fixed tissues

Embryos were mounted individually under a coverslip supported by a tape bridge on either side. This flattened the embryos sufficiently so that all cells were roughly in the same z-plane. *In-situ* HCR stained embryos and immunostained embryos were imaged on an inverted SP8 Confocal Microscope (Leica Systems), with either a 20x 0.75NA air objective, 40X 1.3NA oil-immersion objective, or 63x 1.4NA oil-immersion objective. Either a PMT or HyD detector was used alongside a 405/488/546/594/647nm laser line. Image stacks of various Z separations were captured using the Leica Application Suite X Software.

### Fly strains

We used *yw^67^* as control. Mutant alleles were: *trn^28.4^* for *tartan* (Chang et al., 1993), *ftz^11^* for *fushi-tarazu, 18w^k02701^* for *toll-2* (Yagi et al., 2010) and *comm^E39^* for *commissureless* (Georgiou and Tear, 2002). Transgenes were: *Gap43mCherry* (Martin et al., 2009) to label cell membranes, *sqh^EGFP.29B^* (Proag et al., 2019) to label Myosin II, EnVT15159-peve-MS2-lacZ (this work), nos-MCP-eGFP on II (Garcia et al., 2013), armGal4 (Sanson et al., 1996) and UASwg (Lawrence et al., 1996).

### dsRNA generation

Design of dsRNA was based upon the Heidelberg 2 (BKN) library (Horn and Boutros, 2013). First, to generate transcription templates for production of dsRNA, a PCR was undertaken on *yw^67^* fly gDNA using a Q5 polymerase master mix (NEB) and the following primer pair (preceded by the T7 promoter sequence: 5’-TAATACGACTCACTATAGGG-3’):

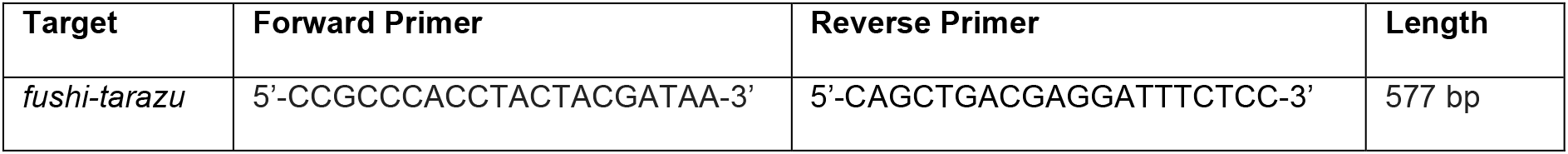

ssRNA was transcribed directly from the PCR amplicon product in a reverse transcription reaction using a HiScribe T7 polymerase (NEB). The DNA template was then removed through treatment with DNase1. ssRNA was annealed to form dsRNA through addition of 0.5M EDTA, 10% SDS and 3M NaCl, boiling the mixture then cooling to RT naturally. Annealed dsRNA was purified through a standard Phenol:Chloroform:IAA 25:24:1 extraction and precipitated from solution by adding of ethanol and ammonium acetate. The isolated dsRNA pellet was washed multiple times in 70% EtOH, air dried, and resuspended in injection buffer (0.1mM sodium phosphate buffer, 5mM KCl). The dsRNA was injected into pre-cellularised embryos at a concentration of 1.7μg/μL, as measured by nanodrop.

### EnVT15159-MS2 generation

The Stark Lab fly enhancer library was used to identify a small region of the *engrailed* enhancer that accurately recapitulates expression at germ-band extended stages of embryogenesis. Tile ID VT15159 contained a 2099bp region of DNA that neighbours the *engrailed* gene and drives *LacZ* in an *engrailed* pattern. The 2099bp region (EnVT15159) was PCR amplified from purified *yw^67^* gDNA using Q5 DNA polymerase MasterMix (NEB) and the following primers:

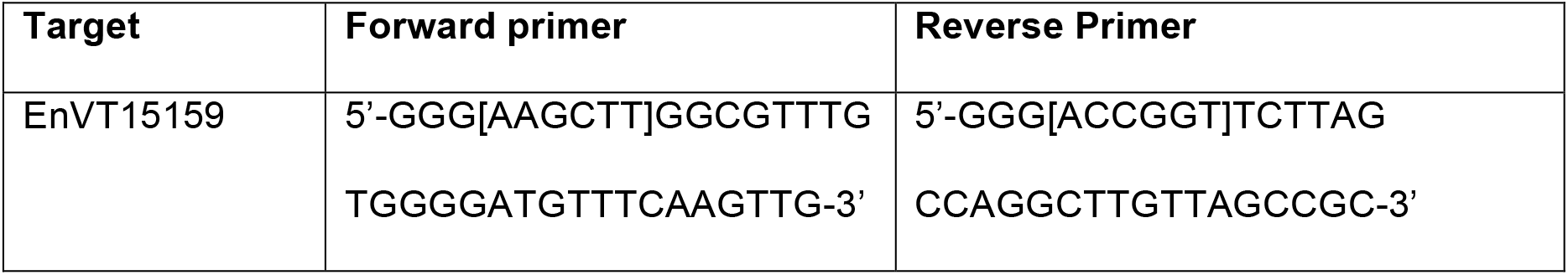

Square brackets indicate HindIII (Fwd primer) and AgeI (Rev primer) restriction enzyme cut sites. Primers were designed so the restriction cut site was preceded by 3 Guanine bases.

2μL of the PCR product was ran on a 1% agarose gel to confirm successful amplification of the EnVT15159 region. The PCR product was cleaned using the Qiaquick PCR clean-up kit (Qiagen). The EnVT15159 PCR product was then digested using HindIII and AgeI high fidelity restriction enzymes (NEB) in Cutsmart buffer (NEB).

The digested EnVT15159 region was cloned into *pattB-w+-pEve-24xMS2-LacZ plasmid* (kind gift of Julia Falo-Sanjuan and Sarah Bray) using the HindIII and *AgeI* sites. The plasmid was transformed into DH5-alpha library-efficiency competent cells (Invitrogen) through the standard heat shock protocol. Transformed colonies (displaying ampicillin resistance) were picked, grown into 50ml of culture and isolated via a MaxiPrep kit (Qiagen). Plasmid fingerprinting was undertaken using Bbs1 and EcoRV restriction enzymes (NEB) to confirm the EnVT15159 product had been inserted into the plasmid in the correct orientation. We injected the final plasmid construct *pattB-w+-EnVT15159-pEve-24xMS2-LacZ* into yw, M(eGFP, vas-int, dmRFP)ZH-2A;; M(attP)ZH-86Fb flies (sourced from Genetics Department Fly Facility, University of Cambridge). The construct was inserted, by phiC31 mediated integration into the attP-86Fb site on the third chromosome (86F8). F1 Transgenic flies were identified through the presence of w+. Crosses were undertaken to generate yw;;EnVT15159-MS2 flies that were homozygous viable and established as a stable stock.

### Immunostaining and antibodies

Embryos were fast fixed at the interface between 37% formaldehyde and 100% heptane for 8 minutes then washed thoroughly in PBS-TX. The vitelline membrane was manually removed through physical manipulation with a tungsten needle. Embryos were blocked in PBS-TX-BSA for 30mins at RT. Embryos were incubated with primary antibodies in blocking solution overnight at 4°C. Excess antibody was removed by washing embryos thoroughly in PBS-TX. Embryos were incubated with secondary antibodies in blocking solution for 1 hour at RT. Excess antibodies were removed by washing thoroughly in PBS-TX. Stained embryos were stored in Vectashield (Vectorlabs) until mounted.

Primary antibodies used were: mouse anti-phospho-Tyrosine (Cell signaling #9411; 1:100), rabbit anti-Engrailed (Santa Cruz D300), chick anti-Beta-gal (Abcam ab9361), rabbit anti-Tartan (Chang et al., 1993)(kind gift of Shigeo Hayashi), rabbit anti-comm (Tear et al., 1996)(kind gift of Guy Tear).

Secondary antibodies conjugated to fluorescent dyes were obtained from Jackson ImmunoResearch Laboratories, Invitrogen and Life Technologies. Streptavidin with Alexa Fluor 405 conjugate was from ThermoFisher Scientific.

### Live imaging

Dechorionated embryos were mounted using an adapted hanging drop methodology (Reed et al., 2009). Briefly, a 22×64mm coverslip (#1) was attached to a rectangular metal microscope slide frame (Leica) using magictape (Scotch). Live embryos freely suspended in Voltalef (PCTFE - H10S - Arkema) were positioned with their ventral side towards the coverslip. The frame and coverslip were quickly inverted. The ventral side of the embryo remains in contact with the coverslip. Embryos were imaged under a 40x oil objective lens (NA of 1.3) on a Nikon Eclipse E1000 microscope with a Yokogawa CSU10 spinning disk head and a Hamamatsu EM-CCD camera. Embryos were illuminated using a Spectral Applied Research LMM2 laser module (491 nm and 561 nm excitation). Images were captured using Volocity Acquisition Software (PerkinElmer) 32 Z-slices with a 1μm separation were obtained at each time point. Embryos were imaged every 30 s from late stage 5 for 100 min. Movies were recorded at 20.5 ± 1°C, measured with a high-resolution thermometer (Checktemp1). To check that embryos survived the imaging process to the end of embryogenesis, embryos were allowed to develop on the imaging insert to hatching in a humidified box. For mutants that are embryonic lethal, the cuticle of embryos was prepared using standard methods to check their phenotype. Occasional movies acquired for embryos that did not hatch or did not make a cuticle at the end of embryogenesis were discarded.

### Cell tracking analysis

Cell Tracking, spatiotemporal movie synchronization, domain strain rate calculations, cell selection criteria and contoured heat map generation were performed as in (Tetley et al., 2016).

### Defining PSB interfaces and cell types

Tissue domains were defined in individual tracked movies by examining the position of EnVT15159-MS2 / MCP-GFP transcriptional dots. To assign dots to cells, cell tracking was undertaken using the custom oTracks software (Blanchard et al., 2009) on movies of embryos containing EnVT15159-MS2, MCP-GFP and Gap43-mCherry. First, the MS2-MCP signal was processed, and a pixel intensity threshold applied to identify dots in an automated manner. Next, the Gap43-mCherry signal was processed and a blanket correction applied to uncurve the 3D surface of the embryo. A few slices of the Gap43-mCherry signal, just under the apical surface of cells, were maximum intensity projected so cell outlines were clear and individual cells could be segmented. Segmented cells were tracked back and forth through time and each cell was marked with a unique identity. Fluorescent transcription dots (resulting from MS2-MCP binding) were also tracked back and forth through time and each dot was assigned to a corresponding cell. Based upon the assignment of dots, *engrailed* expressing cells were identified and cells could then be classified into parasegments meaning PSB interfaces could also be identified. Because cells were tracked over time, these classifications of parasegment identity could be automatically tracked through time to define the same groups of cells at all earlier and later time points.

### Quantifying interface co-alignment

Interface orientations, relative to the embryonic axes, were calculated for PSB, −1 and +1 in all movies. All distributions of interface orientations (from 0, parallel to the AP embryonic axis, to 180°) were reflected around 90°, producing distributions from 0°, AP-aligned, to 90°, DV-aligned. As a measure of co-alignment, the proportion of interfaces oriented between 60 and 90° relative to the AP axis was plotted over time, from −10 to 50 min. Cumulative frequencies were calculated for each reflected distribution of interface orientations. Two-sample Kolmogorov-Smirnov tests on the cumulative frequency distributions of interface orientation were used to compare treatments / genotypes.

### Cuticle preparation

Dechorionated embryos were transferred to a 50:50 mix of Hoyer’s medium and lactic acid and mounted under a 22×32mm coverslip. A permanent marker was used to draw black dots onto the surface of the coverslip to help locate embryos for microscopy. The slide was baked overnight at 62°C with a weight on top of the cuticle to prevent air bubbles forming. Cuticle preps were imaged using darkfield or phase-contrast microscopy.

### Embryo injections

Adult flies were kept at 25°C in a cage with an apple juice agar plate. Embryos were collected from the plate following a 30-minute laying period and were dechorionated. Approximately 20 embryos (for dsRNAi experiments) and 100 embryos (for transgenic injections) were aligned on a block of agar and transferred to a coverslip using a thin layer of heptane glue. If the injected embryos were for live imaging, embryos would be aligned with their ventral side facing the glue and coverslip. Embryos were desiccated in a jar of silica beads for 10 to 12 minutes before being covered with a thin layer of Voltalef (PCTFE H10S, Arkema). A brightfield microscope (Nikon), microinjection platform (Leica), and a pulled glass needle were used to inject the embryos through their posterior end. For RNA interference experiments, the expulsion of dsRNA was aimed for the centre of the embryo. For the generation of transgenic flies, plasmid solution was injected at the posterior end of the embryo (where the future pole cells form). Unfertilised, damaged, or old embryos were destroyed with forceps. Slides of injected embryos were placed in a 50mm petri dish at 18°C until the correct stage of development.

## Supplementary figures legends

**Supplemental Figure 1:**
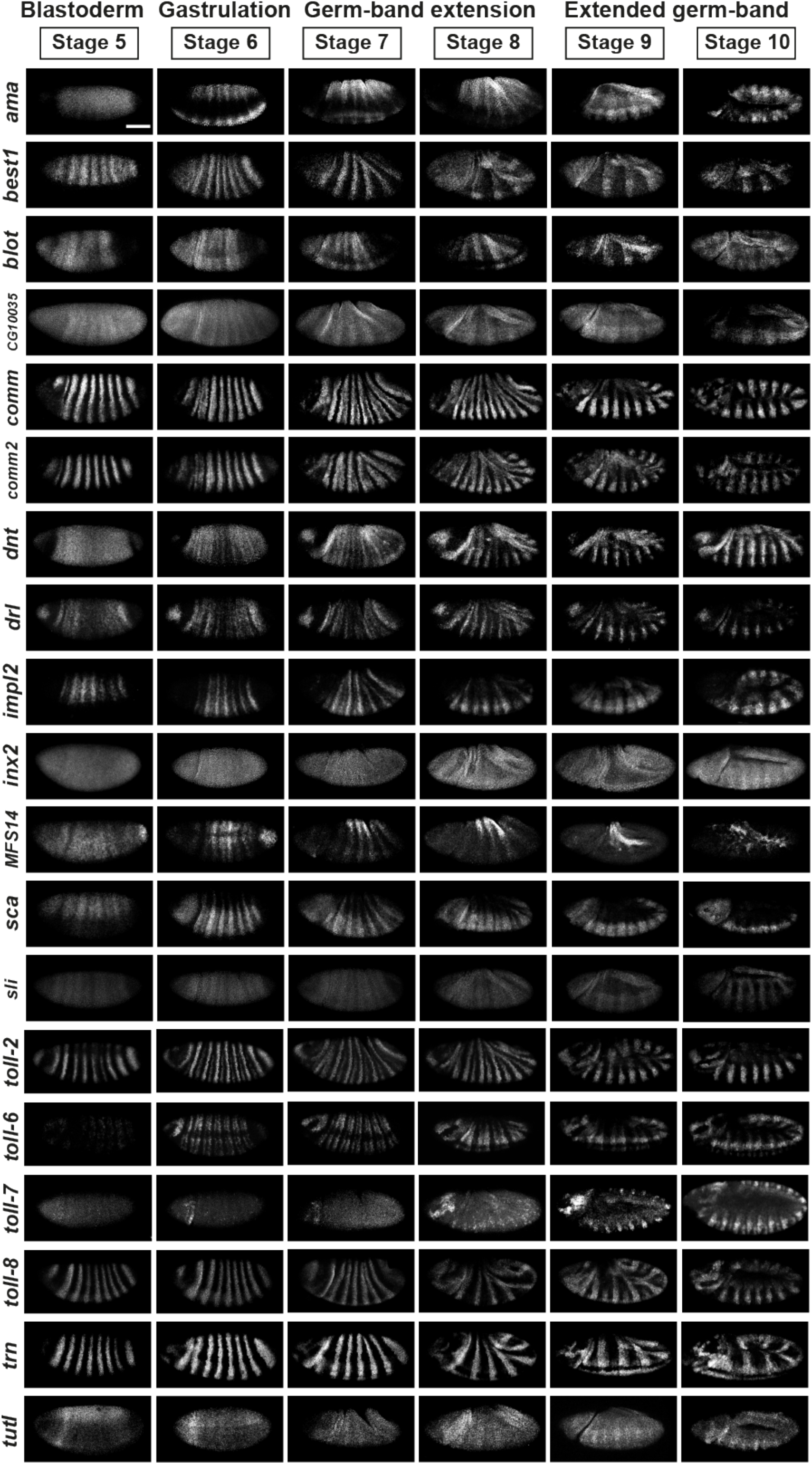
*In situ* HCR for candidate genes with antero-posterior striped expression. HCR was performed for each gene in *yw^67^ Drosophila* embryos between stages 5 to 10 of embryogenesis. The 19 genes included in this figure were classified as having a striped expression pattern at some point during stages 5-10. Some genes, such as *tutl*, subtle stripes are only clearly visible at a subset of the stages surveyed (stages 9-10), whereas other genes, such as *trn*, have clear expression stripes visible at all stages. Scale bar = 100μm.

**Supplemental Figure 2:**
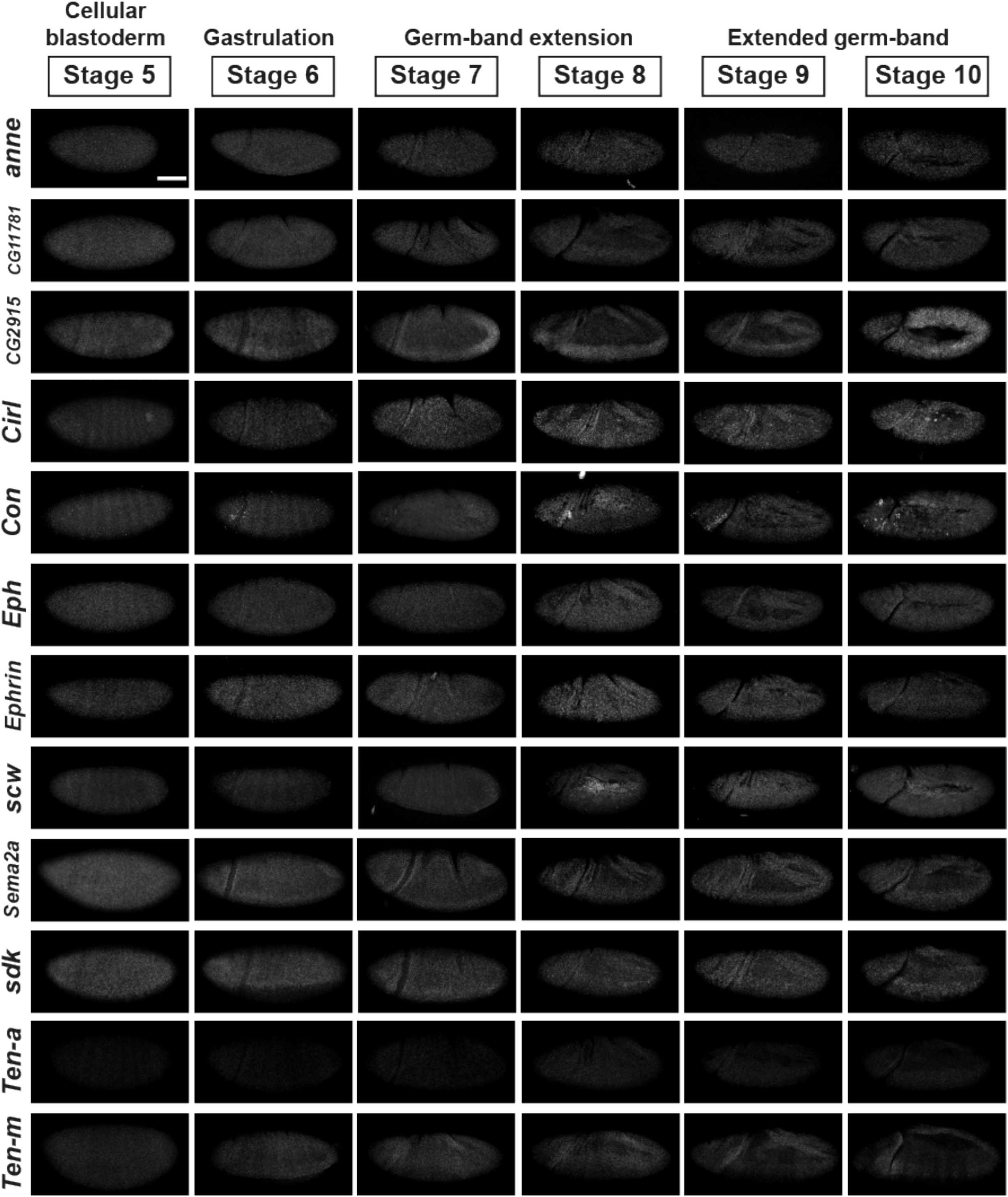
*In situ* HCR for candidate genes without AP striped expression pattern. HCR was performed for each gene in *yw^67^ Drosophila* embryos between stages 5 to 10 of embryogenesis and the 12 genes included here were found not to have any AP striped expression.

**Supplemental Figure 3:**
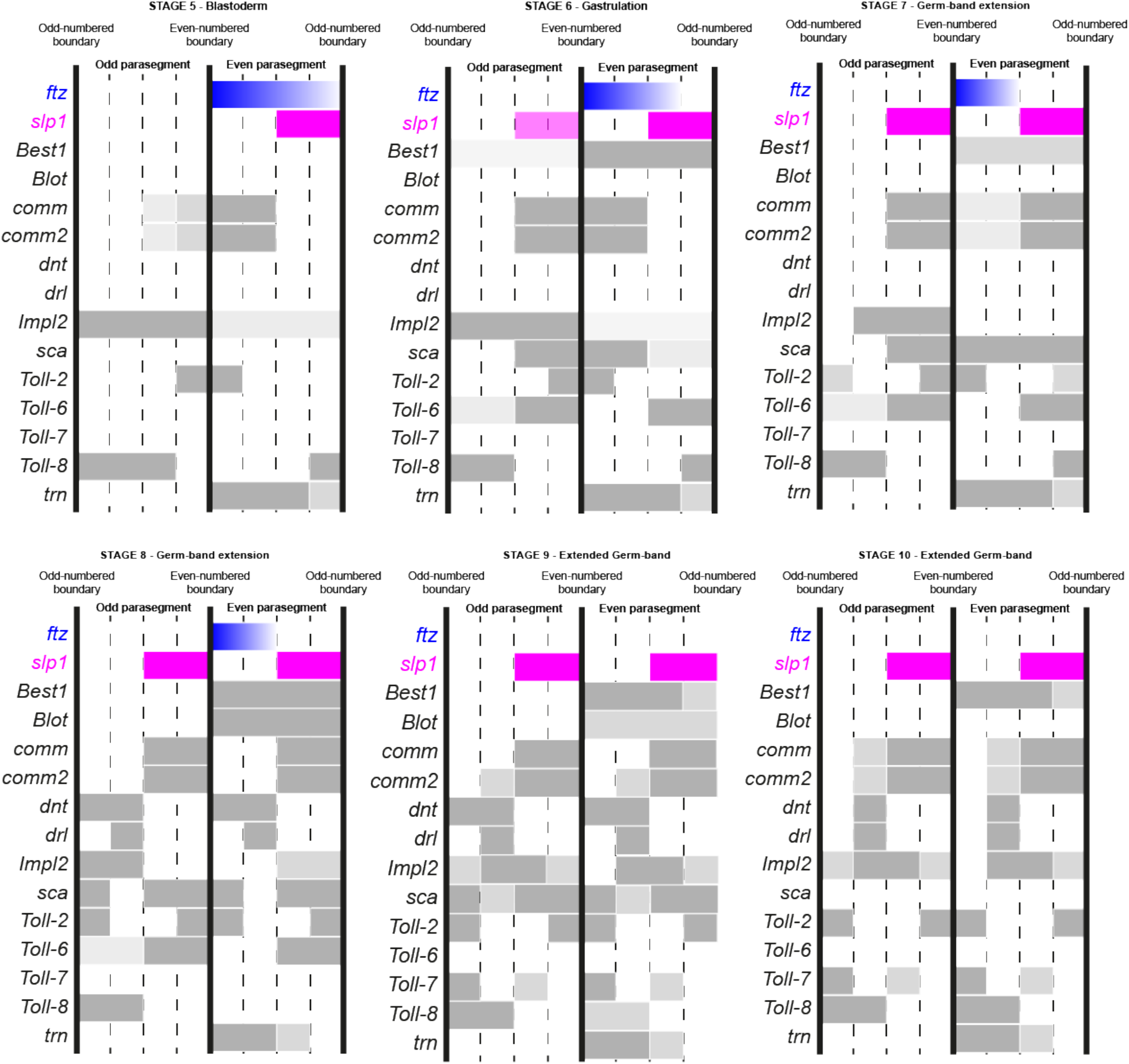
Map of expression patterns of candidate genes relative to the PSBs. The maps are drawn based on gene expression patterns revealed by HCR for each candidate genes in combination with the parasegmental boundary markers *ftz* and *slp1*. HCR was performed in *yw^67^ Drosophila* embryos between stages 5 to 10 of embryogenesis. Out of 19 candidates with AP striped expression, the 13 genes included here showed expression patterns that were clear enough to enable mapping relative to the PSBs. Note that for Tartan, expression at stages 9 and 10 is not continuous along DV anymore.

**Supplemental Figure 4:**
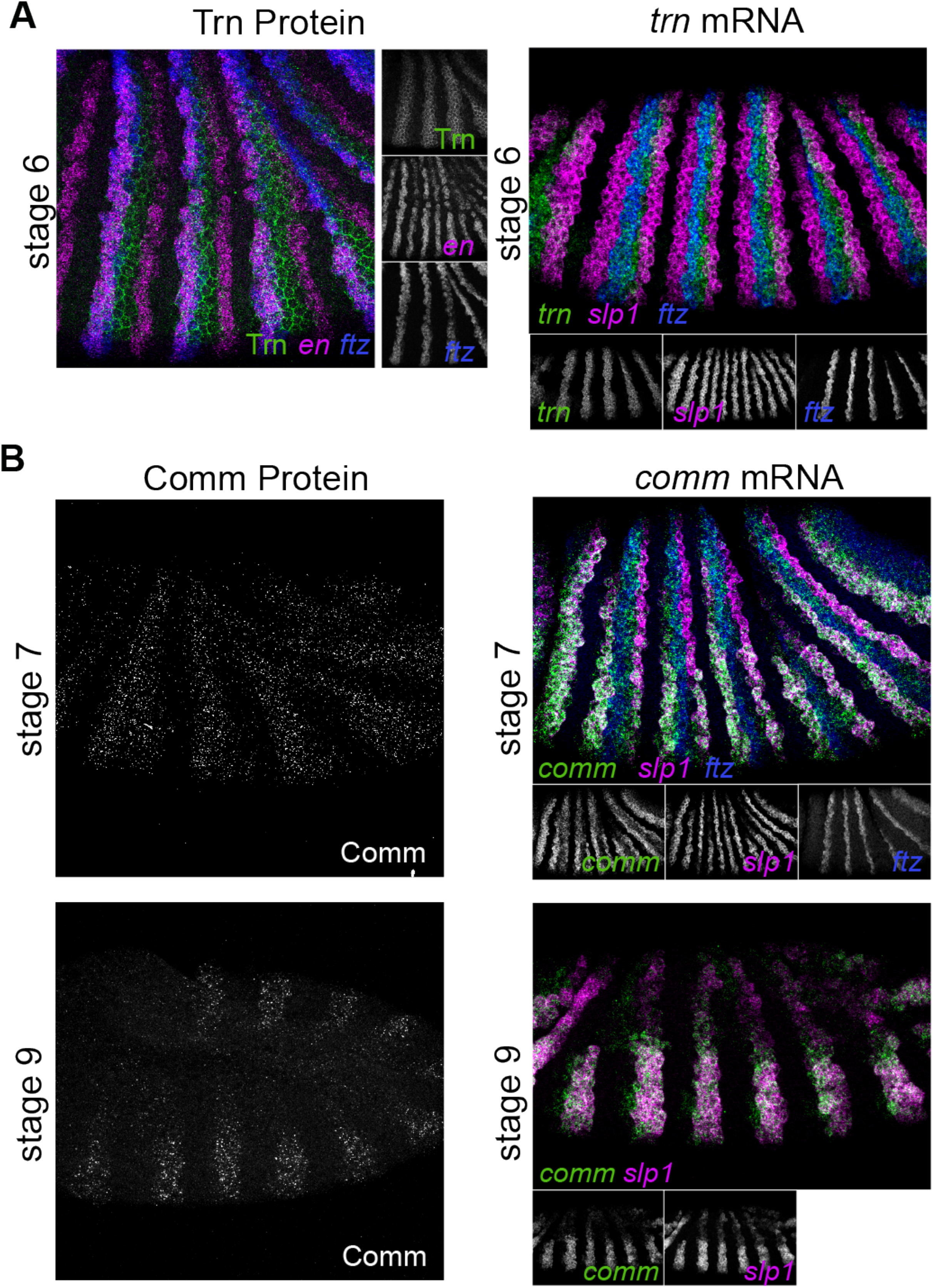
Comparison of mRNA and protein patterns for candidate genes *tartan* and *comm*. A) Left panel shows an immunostaining against the Tartan protein in combination with HCR for *en* and *ftz*. Right panel show HCR for *tartan*, in combination with *slp1* and *ftz*. Both images show a stage 7 *yw^67^* embryo. For the immunostain, maximum intensity projection are shown for the full z-depth of the *engrailed* and *fushi-tarazu* channels and for 2μm of Tartan signal, just below the apical surface of cells. Tartan protein localises to the membrane of cells located within even-numbered parasegments and is absent from cells within odd-numbered parasegments at stages 6-8 of embryogenesis. (Scale bar = 50μm). B) Left panel show immunostainings against Comm protein and right panel HCR for comm, in combination with *slp1* and *ftz* markers. Images of stage 8 and 9 *yw^67^* embryos show that *comm* patterns for both mRNA and protein undergo a doubling of periodicity (see also Sup. Fig 3).

**Supplemental Figure 5:**
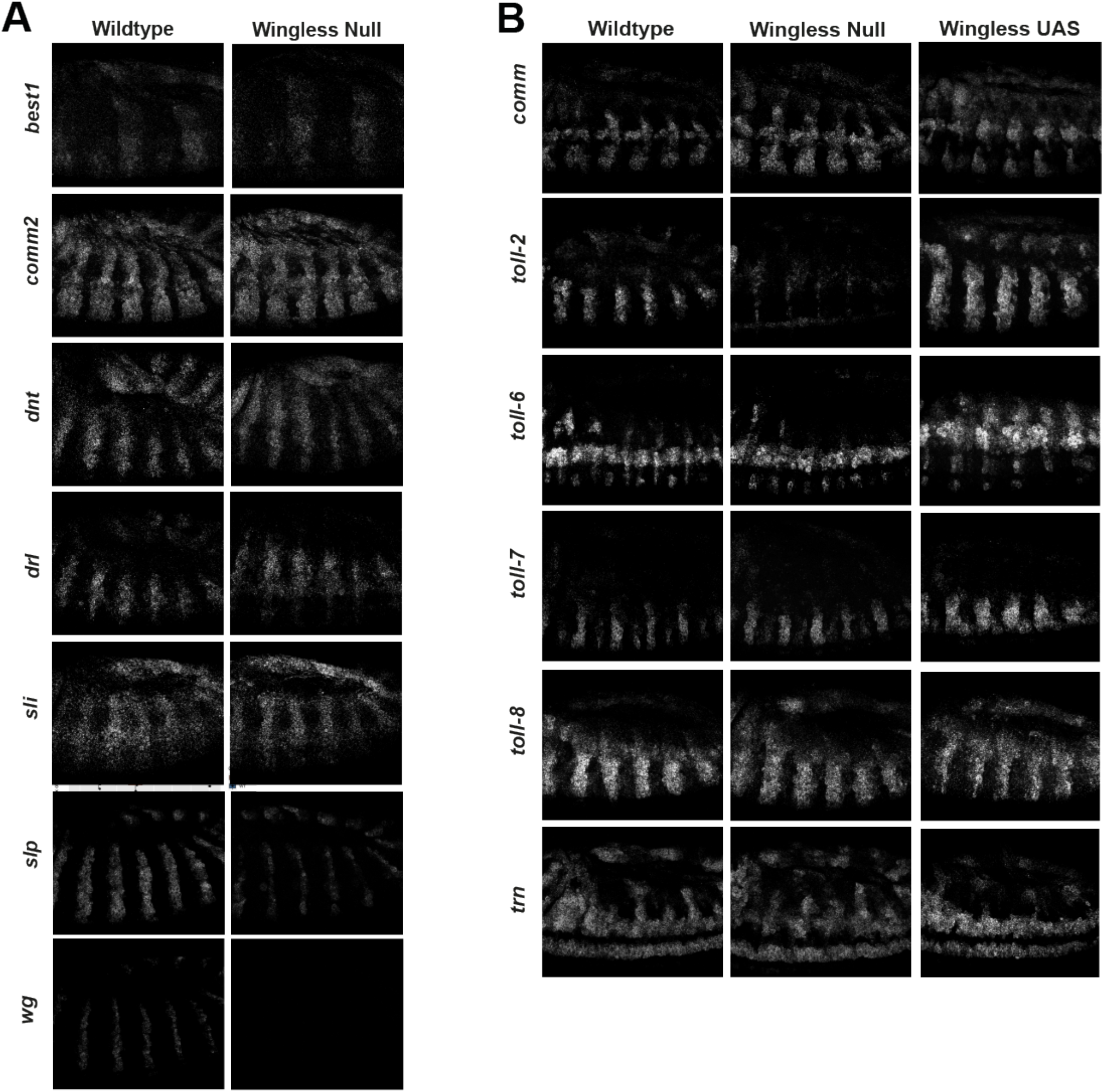
*In situ* HCR to identify candidate genes regulated by Wingless signalling. A) HCR for a subset of candidate genes in WT and *wg^cx4^* mutant embryos, alongside *wg* and *slp1* controls. B) HCR for an additional subset of candidate genes in WT, *wg^cx4^* mutant embryos and in *armGal4/UASwg* embryos expressing Wg ubiquitously. Out of all the genes tested, only *toll-2* shows robust change in HCR, with loss of signal in *wg^cx4^* mutant and increase in *armGal4/UASwg* embryos.

**Supplemental Figure 6:**
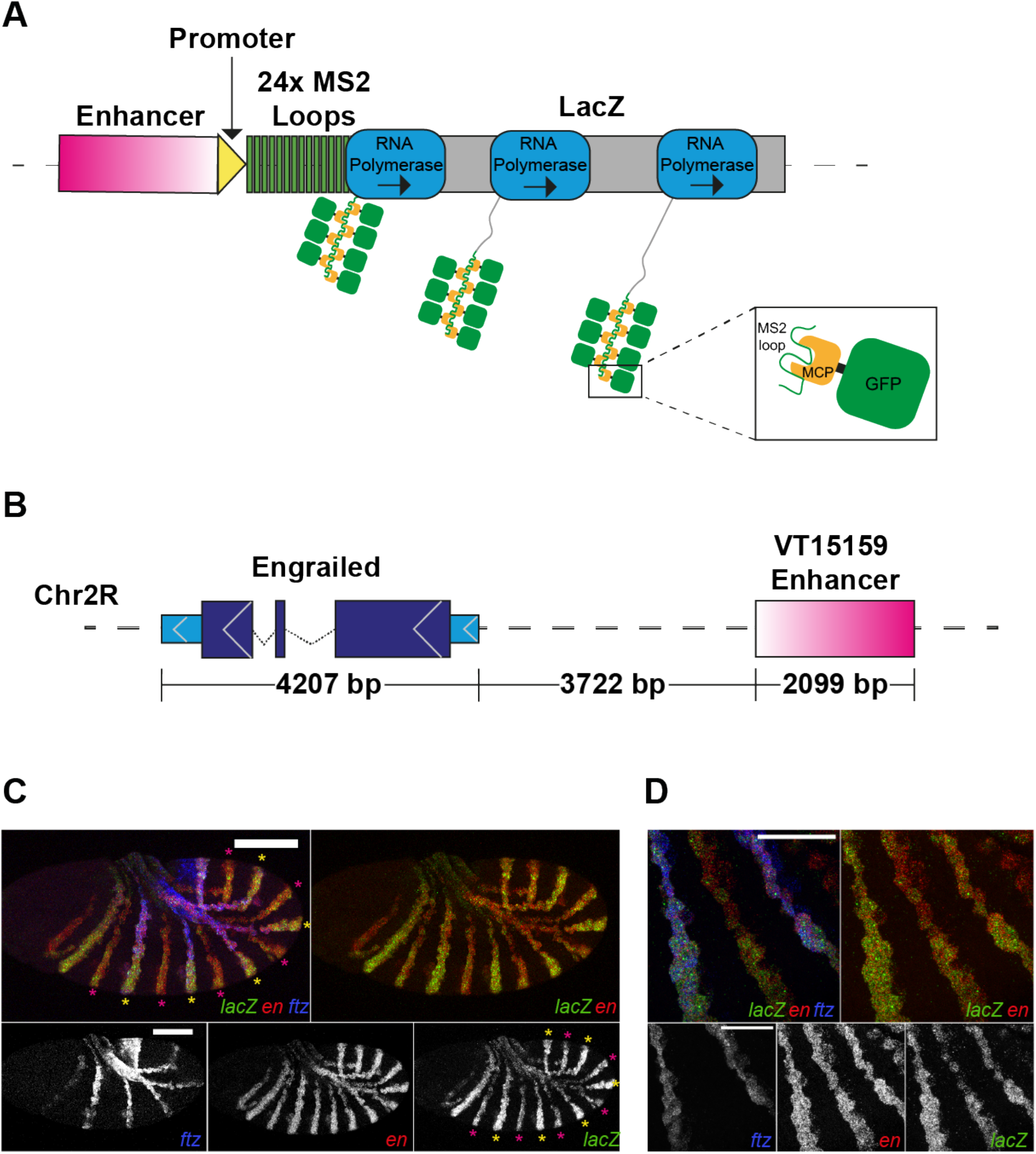
MS2-MCP system to image nascent mRNA transcripts from an *engrailed* enhancer in live embryos. **A)** Diagram showing the MS2-MCP system. The gene of interest’s enhancer region is cloned upstream of a strong promoter that initiates transcription. RNA polymerase transcribes the 24 MS2 stem loops followed by *lacZ*. The resulting mRNAs contains 24 copies of the MS2 loops that are bound by the MCP protein fused to GFP (box). The concentration of multiple MCP-GFP/MS2 loop complexes causes a spatially localised fluorescence that is detected at the locus as a fluorescent dot. Multiple transcripts are produced from a single locus simultaneously increasing the brightness of the fluorescent dot. *lacZ* transcription is accessory to the core system but prolongs the time the nascent mRNA stays within vicinity of the locus, making the fluorescent dot brighter. B) Gene diagram showing the position of the VT15159 enhancer region, which is located 3722bp upstream of the *engrailed* gene (adapted from the NCBI genome data viewer). C) HCR of *ftz* (blue), *engrailed* (red) and *lacZ* (green) in embryos transgenic for EnVT15159-MS2. *lacZ* expression from EnVT15159-MS2 coincides with endogenous *engrailed* expression demonstrating that EnVT15159 recapitulates endogenous *engrailed* expression during early embryogenesis. Note that brighter alternate *lacZ* stripes overlap with *ftz* expression, indicating differential patterning by the VT15159 between odd and even-numbered parasegments, with the brighter stripes abutting even-numbered PSBs. (20x magnification. Scale bar = 100μm) D) Close-up to show the coincidence between endogenous engrailed expression and *lacZ* expression from EnVT15159. The brighter *lacZ* stripes overlap with *ftz* expression and abut even-numbered PSBs. (63x magnification and 2x zoom. Scale bar = 50μm)

**Supplemental Figure 7:**
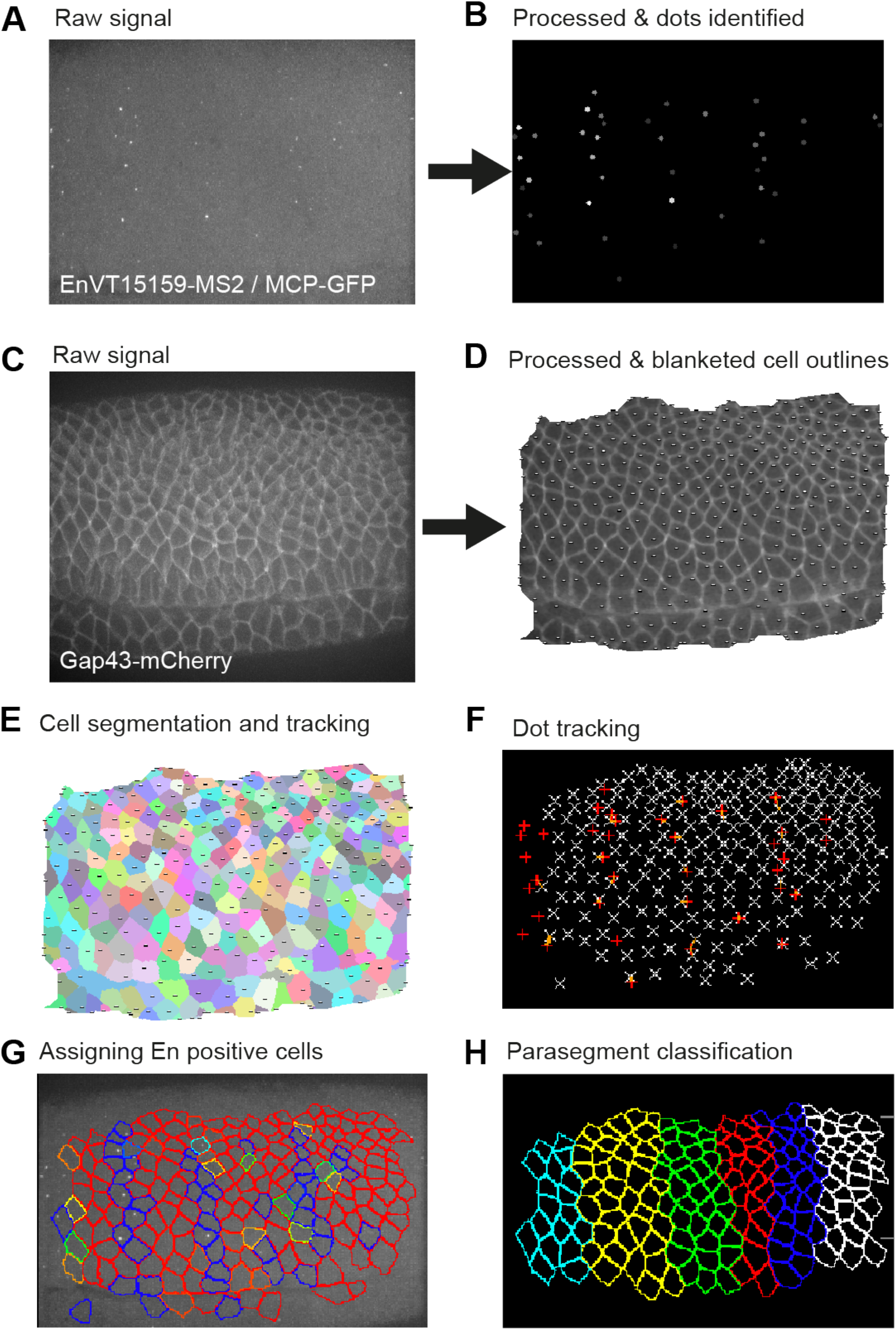
Image processing and cell tracking pipeline to identify parasegmental boundaries in live embryos. A,B) Raw EnVT15159-MS2-MCP-GFP signal is processed in the custom software oTracks. A median background subtraction of 10 is applied. Fluorescent dots are identified in an automated manner by applying a pixel intensity threshold. C,D) Raw Gap43-mCherry signal is processed within oTracks. Median smoothing of 1 is applied and the corners of the image are brightened to correct for microscope artefacts. A blanketing operation is applied to correct for the 3D curvature of the embryonic volume. A 2μm z-plane is projected from the embryonic volume onto a flat 2D surface for image segmentation. E) oTracks segments each cell based on the Gap43-mCherry signal using a watershed algorithm. Each cell is tracked back and forth through frames and connected in time. F) Identified dots are tracked back and forth through time in oTracks. Dots are assigned a probability of belonging to each cell by determining the proximity of each dot to each cell during the lifetime of each dot. G) The probability of a cell being assigned to a dot can be displayed as a color (red = low probability, blue = high probability). Cells with a high probability of containing a dot form AP stripes throughout the embryo similar to the *engrailed* expression pattern. H) Based upon the color coding in G, parasegment identities can be assigned to each cell and the location of PSBs can be identified.

**Supplemental Figure 8:**
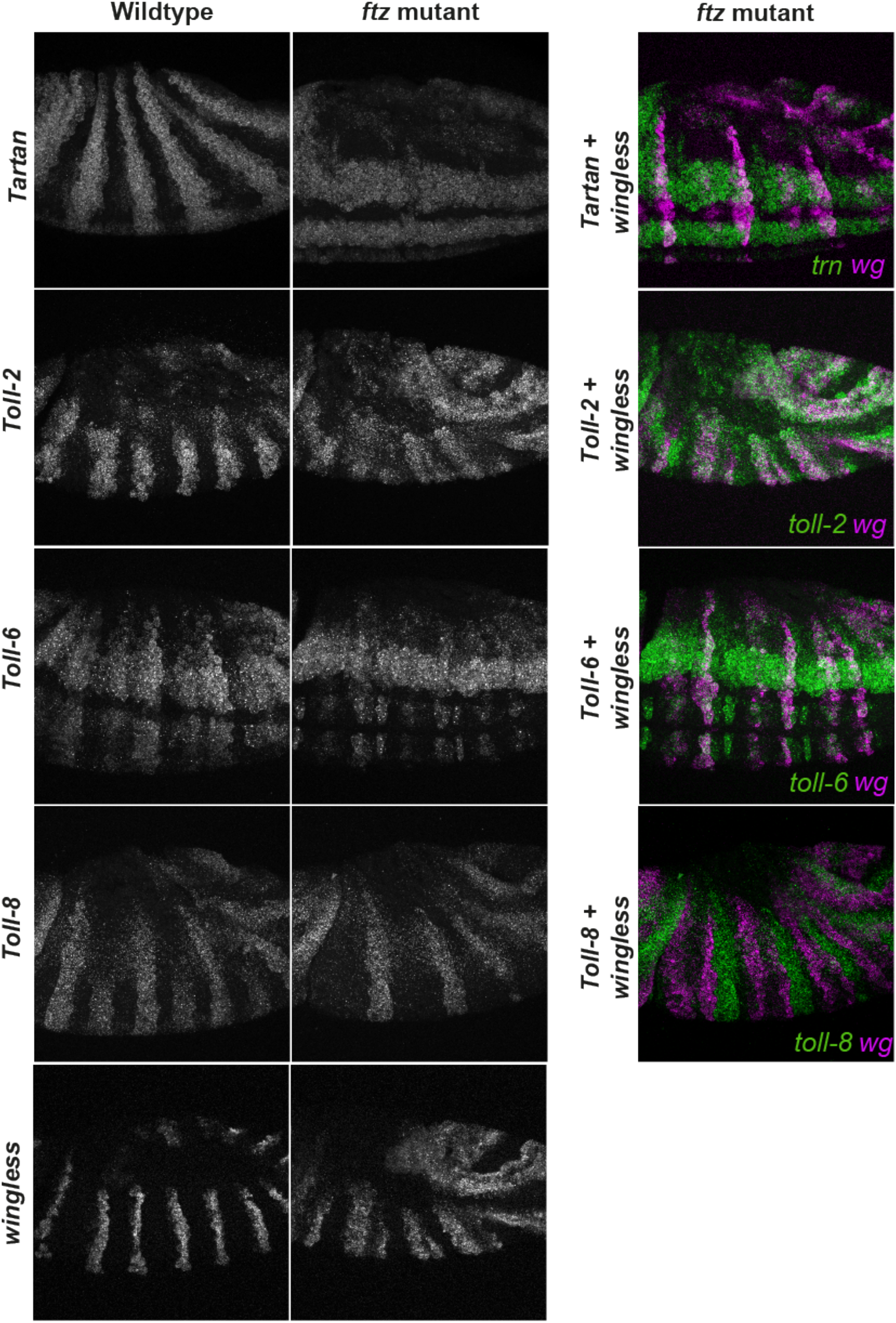
Expression of LRR transmembrane receptors in *ftz* mutants. HCR of *tartan, toll-2, toll-6* and *toll-8* on their own (left panels) and in combination with *wg* as a marker (right panels), in wildtype and *ftz* mutant embryos. *ftz* mutant embryos are homozygous for the *ftz^11^* null allele while wildtype embryos are heterozygous. *ftz* mutant embryos were identified based on the changes in *wg* expression pattern.

**Supplemental Figure 9:**
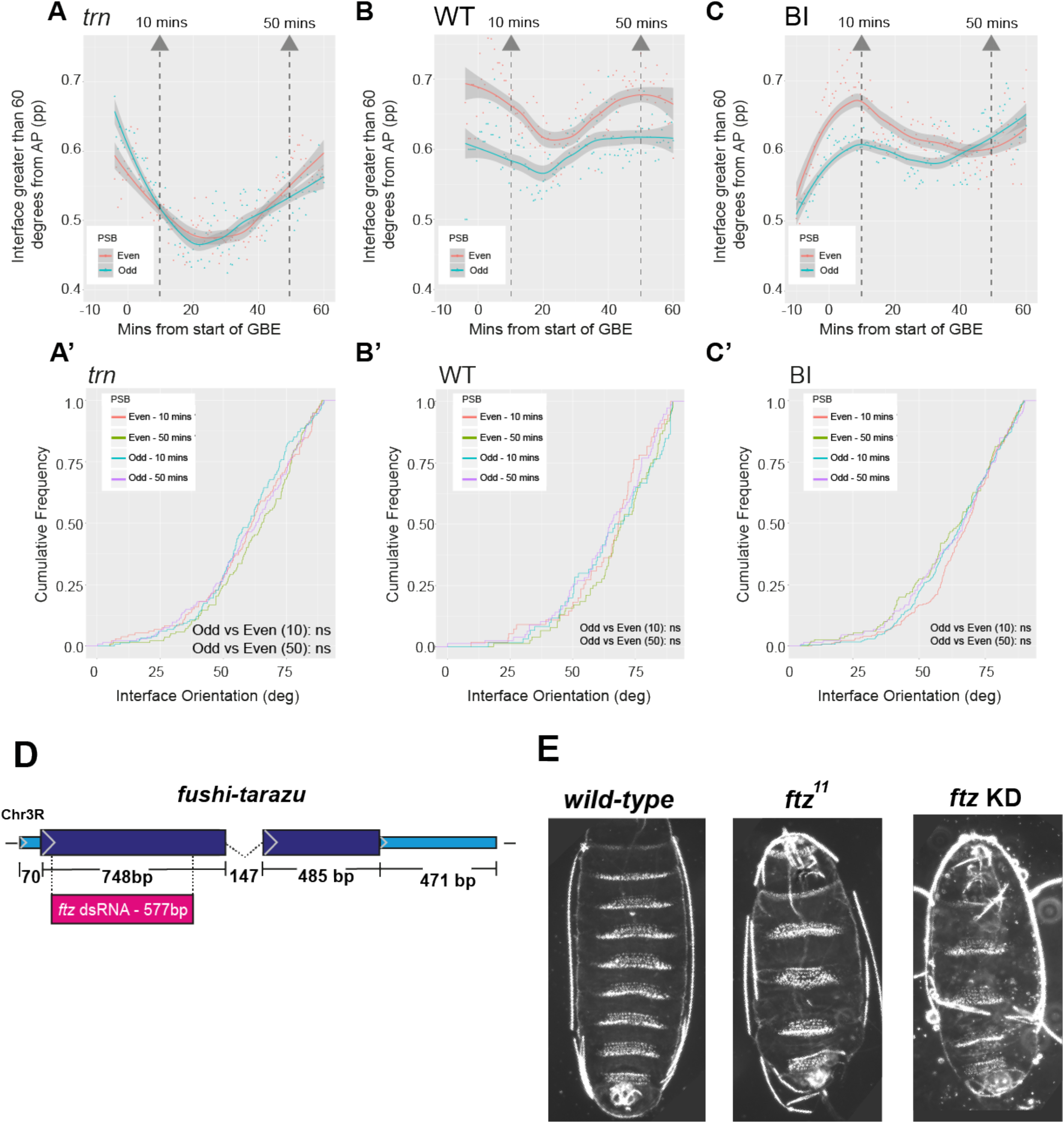
Boundary straightness comparisons between even and odd-numbered PSBs and *ftz* dsRNA knockdowns. A-C) Plots showing the proportion of odd and even-numbered PSB boundary interfaces that are greater than 60 degrees from AP in 3 *tartan* mutant (A,A’), 3 wildtype (B,B’) and 4 buffer injected embryos (C, C’). Odd and even-numbered PSBs are distinguished by the differential expression of EnVT15159-MS2 transcriptional dots. A loess curve (span 0.75) has been fitted to the data. Statistical comparison is a Kolmogorov-Smirnov non-parametric test undertaken on the cumulative frequencies of interface angles at 10 and 50 minutes (shown in A’-C’). D) *ftz* gene model (adapted from NCBI genome viewer) showing the dsRNA target sequence in exon 1 of *ftz* mRNA. E) Dark field microscopy images of cuticle preps from wildtype, *ftz^11^* null and ftz dsRNA knockdown embryos. The ftz KD phenocopies the *ftz* null mutant, displaying the expected pair-rule phenotype.

## References

Aliee, M., Roper, J. C., Landsberg, K. P., Pentzold, C., Widmann, T. J., Julicher, F. and Dahmann, C. (2012). Physical mechanisms shaping the Drosophila dorsoventral compartment boundary. Curr Biol 22, 967–976.

Amack, J. D. and Manning, M. L. (2012). Knowing the boundaries: extending the differential adhesion hypothesis in embryonic cell sorting. Science 338, 212–215.

Bertet, C., Sulak, L. and Lecuit, T. (2004). Myosin-dependent junction remodelling controls planar cell intercalation and axis elongation. Nature 429, 667–671.

Bielmeier, C., Alt, S., Weichselberger, V., La Fortezza, M., Harz, H., Julicher, F., Salbreux, G. and Classen, A. K. (2016). Interface Contractility between Differently Fated Cells Drives Cell Elimination and Cyst Formation. Curr Biol 26, 563–574.

Blanchard, G. B., Kabla, A. J., Schultz, N. L., Butler, L. C., Sanson, B., Gorfinkiel, N., Mahadevan, L. and Adams, R. J. (2009). Tissue tectonics: morphogenetic strain rates, cell shape change and intercalation. Nat Methods 6, 458–464.

Bosveld, F., Guirao, B., Wang, Z., Riviere, M., Bonnet, I., Graner, F. and Bellaiche, Y. (2016). Modulation of junction tension by tumor suppressors and proto-oncogenes regulates cell-cell contacts. Development 143, 623–634.

Butler, L. C., Blanchard, G. B., Kabla, A. J., Lawrence, N. J., Welchman, D. P., Mahadevan, L., Adams, R. J. and Sanson, B. (2009). Cell shape changes indicate a role for extrinsic tensile forces in Drosophila germ-band extension. Nat Cell Biol 11, 859–864.

Calzolari, S., Terriente, J. and Pujades, C. (2014). Cell segregation in the vertebrate hindbrain relies on actomyosin cables located at the interhombomeric boundaries. EMBO J 33, 686–701.

Canty, L., Zarour, E., Kashkooli, L., Francois, P. and Fagotto, F. (2017). Sorting at embryonic boundaries requires high heterotypic interfacial tension. Nature communications 8, 157.

Chang, Z., Price, B. D., Bockheim, S., Boedigheimer, M. J., Smith, R. and Laughon, A. (1993). Molecular and genetic characterization of the Drosophila tartan gene. Dev Biol 160, 315–332.

Choi, H. M. T., Schwarzkopf, M., Fornace, M. E., Acharya, A., Artavanis, G., Stegmaier, J., Cunha, A. and Pierce, N. A. (2018). Third-generation in situ hybridization chain reaction: multiplexed, quantitative, sensitive, versatile, robust. Development 145.

Collinet, C. and Lecuit, T. (2021). Programmed and self-organized flow of information during morphogenesis. Nat Rev Mol Cell Biol 22, 245–265.

Dahmann, C., Oates, A. C. and Brand, M. (2011). Boundary formation and maintenance in tissue development. Nat Rev Genet 12, 43–55.

DiNardo, S., Sher, E., Heemskerk-Jongens, J., Kassis, J. A. and O’Farrell, P. H. (1988). Two-tiered regulation of spatially patterned engrailed gene expression during Drosophila embryogenesis. Nature 332, 604–609.

Fagotto, F. (2020a). Cell sorting at embryonic boundaries. Semin Cell Dev Biol 107, 126–129.

Fagotto, F. (2020b). Tissue segregation in the early vertebrate embryo. Semin Cell Dev Biol 107, 130–146.

Fernandez-Gonzalez, R., Simoes Sde, M., Roper, J. C., Eaton, S. and Zallen, J. A. (2009). Myosin II dynamics are regulated by tension in intercalating cells. Dev Cell 17, 736–743.

Florence, B., Guichet, A., Ephrussi, A. and Laughon, A. (1997). Ftz-F1 is a cofactor in Ftz activation of the Drosophila engrailed gene. Development 124, 839–847.

Garcia De Las Bayonas, A., Philippe, J. M., Lellouch, A. C. and Lecuit, T. (2019). Distinct RhoGEFs Activate Apical and Junctional Contractility under Control of G Proteins during Epithelial Morphogenesis. Curr Biol 29, 3370–3385 e3377.

Garcia, H. G., Tikhonov, M., Lin, A. and Gregor, T. (2013). Quantitative imaging of transcription in living Drosophila embryos links polymerase activity to patterning. Curr Biol 23, 2140–2145.

Georgiou, M. and Tear, G. (2002). Commissureless is required both in commissural neurones and midline cells for axon guidance across the midline. Development 129, 2947–2956.

Horn, T. and Boutros, M. (2013). Design of RNAi reagents for invertebrate model organisms and human disease vectors. Methods Mol Biol 942, 315–346.

Howard, K. and Ingham, P. (1986). Regulatory interactions between the segmentation genes fushi tarazu, hairy, and engrailed in the Drosophila blastoderm. Cell 44, 949–957.

Hynes, R. O. and Zhao, Q. (2000). The evolution of cell adhesion. J Cell Biol 150, F89–96.

Ing, B., Shteiman-Kotler, A., Castelli, M., Henry, P., Pak, Y., Stewart, B., Boulianne, G. L. and Rotin, D. (2007). Regulation of Commissureless by the ubiquitin ligase DNedd4 is required for neuromuscular synaptogenesis in Drosophila melanogaster. Mol Cell Biol 27, 481–496.

Irvine, K. D. and Wieschaus, E. (1994). Cell intercalation during Drosophila germband extension and its regulation by pair-rule segmentation genes. Development 120, 827–841.

Justice, E. D., Barnum, S. J. and Kidd, T. (2017). The WAGR syndrome gene PRRG4 is a functional homologue of the commissureless axon guidance gene. PLoS genetics 13, e1006865.

Keleman, K., Rajagopalan, S., Cleppien, D., Teis, D., Paiha, K., Huber, L. A., Technau, G. M. and Dickson, B. J. (2002). Comm sorts robo to control axon guidance at the Drosophila midline. Cell 110, 415–427.

Kerridge, S., Munjal, A., Philippe, J. M., Jha, A., de las Bayonas, A. G., Saurin, A. J. and Lecuit, T. (2016). Modular activation of Rho1 by GPCR signalling imparts polarized myosin II activation during morphogenesis. Nat Cell Biol 18, 261–270.

Landsberg, K. P., Farhadifar, R., Ranft, J., Umetsu, D., Widmann, T. J., Bittig, T., Said, A., Julicher, F. and Dahmann, C. (2009). Increased cell bond tension governs cell sorting at the Drosophila anteroposterior compartment boundary. Curr Biol 19, 1950–1955.

Larsen, C., Bardet, P.-L., Vincent, J.-P. and Alexandre, C. (2008). Specification and positioning of parasegment grooves in Drosophila. Developmental Biology 321, 310–318.

Lavalou, J., Mao, Q., Harmansa, S., Kerridge, S., Lellouch, A. C., Philippe, J. M., Audebert, S., Camoin, L. and Lecuit, T. (2021). Formation of polarized contractile interfaces by self-organized Toll-8/Cirl GPCR asymmetry. Dev Cell 56, 1574–1588 e1577.

Lawrence, P. A., Sanson, B. and Vincent, J. P. (1996). Compartments, wingless and engrailed: patterning the ventral epidermis of Drosophila embryos. Development 122, 4095–4103.

Lye, C. M., Blanchard, G. B., Naylor, H. W., Muresan, L., Huisken, J., Adams, R. J. and Sanson, B. (2015). Mechanical Coupling between Endoderm Invagination and Axis Extension in Drosophila. PLoS Biol 13, e1002292.

Martin, A. C., Kaschube, M. and Wieschaus, E. F. (2009). Pulsed contractions of an actin-myosin network drive apical constriction. Nature 457, 495–499.

Martin, E., Theis, S., Gay, G., Monier, B., Rouviere, C. and Suzanne, M. (2021). Arp2/3-dependent mechanical control of morphogenetic robustness in an inherently challenging environment. Dev Cell 56, 687–701 e687.

Monier, B., Pelissier-Monier, A., Brand, A. H. and Sanson, B. (2010). An actomyosin-based barrier inhibits cell mixing at compartmental boundaries in Drosophila embryos. Nat Cell Biol 12, 60–65.

Monier, B., Pelissier-Monier, A. and Sanson, B. (2011). Establishment and maintenance of compartmental boundaries: role of contractile actomyosin barriers. Cell Mol Life Sci 68, 1897–1910.

Nishimura, T., Honda, H. and Takeichi, M. (2012). Planar cell polarity links axes of spatial dynamics in neural-tube closure. Cell 149, 1084–1097.

Pare, A. C., Naik, P., Shi, J., Mirman, Z., Palmquist, K. H. and Zallen, J. A. (2019). An LRR Receptor-Teneurin System Directs Planar Polarity at Compartment Boundaries. Dev Cell 51, 208–221 e206.

Pare, A. C., Vichas, A., Fincher, C. T., Mirman, Z., Farrell, D. L., Mainieri, A. and Zallen, J. A. (2014). A positional Toll receptor code directs convergent extension in Drosophila. Nature 515, 523–527.

Pare, A. C. and Zallen, J. A. (2020). Cellular, molecular, and biophysical control of epithelial cell intercalation. Curr Top Dev Biol 136, 167–193.

Proag, A., Monier, B. and Suzanne, M. (2019). Physical and functional cell-matrix uncoupling in a developing tissue under tension. Development 146.

Pujades, C. (2020). The multiple functions of hindbrain boundary cells: Tinkering boundaries? Semin Cell Dev Biol 107, 179–189.

Reed, B. H., McMillan, S. C. and Chaudhary, R. (2009). The preparation of Drosophila embryos for live-imaging using the hanging drop protocol. J Vis Exp.

Roy, S., Ernst, J., Kharchenko, P. V., Kheradpour, P., Negre, N., Eaton, M. L., Landolin, J. M., Bristow, C. A., Ma, L., Lin, M. F., et al. (2010). Identification of functional elements and regulatory circuits by Drosophila modENCODE. Science 330, 1787–1797.

Rozbicki, E., Chuai, M., Karjalainen, A. I., Song, F., Sang, H. M., Martin, R., Knolker, H. J., MacDonald, M. P. and Weijer, C. J. (2015). Myosin-II-mediated cell shape changes and cell intercalation contribute to primitive streak formation. Nat Cell Biol 17, 397–408.

Sanson, B., White, P. and Vincent, J. P. (1996). Uncoupling cadherin-based adhesion from wingless signalling in Drosophila. Nature 383, 627–630.

Scarpa, E., Finet, C., Blanchard, G. B. and Sanson, B. (2018). Actomyosin-Driven Tension at Compartmental Boundaries Orients Cell Division Independently of Cell Geometry In Vivo. Dev Cell 47, 727–740 e726.

Sharrock, T. E. and Sanson, B. (2020). Cell sorting and morphogenesis in early Drosophila embryos. Semin Cell Dev Biol 107, 147–160.

Shindo, A. and Wallingford, J. B. (2014). PCP and septins compartmentalize cortical actomyosin to direct collective cell movement. Science 343, 649–652.

Tamada, M., Shi, J., Bourdot, K. S., Supriyatno, S., Palmquist, K. H., Gutierrez-Ruiz, O. L. and Zallen, J. A. (2021). Toll receptors remodel epithelia by directing planar-polarized Src and PI3K activity. Dev Cell 56, 1589–1602 e1589.

Tear, G., Harris, R., Sutaria, S., Kilomanski, K., Goodman, C. S. and Seeger, M. A. (1996). commissureless controls growth cone guidance across the CNS midline in Drosophila and encodes a novel membrane protein. Neuron 16, 501–514.

Tetley, R. J., Blanchard, G. B., Fletcher, A. G., Adams, R. J. and Sanson, B. (2016). Unipolar distributions of junctional Myosin II identify cell stripe boundaries that drive cell intercalation throughout Drosophila axis extension. Elife 5, e12094.

Urbano, J. M., Naylor, H. W., Scarpa, E., Muresan, L. and Sanson, B. (2018). Suppression of epithelial folding at actomyosin-enriched compartment boundaries downstream of Wingless signalling in Drosophila. Development 145.

Wang, J. and Dahmann, C. (2020). Establishing compartment boundaries in Drosophila wing imaginal discs: An interplay between selector genes, signaling pathways and cell mechanics. Semin Cell Dev Biol 107, 161–169.

Yagi, Y., Nishida, Y. and Ip, Y. T. (2010). Functional analysis of Toll-related genes in Drosophila. Development, growth & differentiation 52, 771–783.

Zallen, J. A. and Wieschaus, E. (2004). Patterned gene expression directs bipolar planar polarity in Drosophila. Dev Cell 6, 343–355.

